# Sex Chromosomes and Gonads Shape the Sex-Biased Transcriptomic Landscape in Tlr7-Mediated Demyelination During Aging

**DOI:** 10.1101/2023.09.19.558439

**Authors:** Chloe Lopez-Lee, Lay Kodama, Li Fan, Man Ying Wong, Nessa R. Foxe, Laraib Jiaz, Fangmin Yu, Pearly Ye, Jingjie Zhu, Kendra Norman, Eileen Ruth Torres, Rachel D. Kim, Gergey Alzaem Mousa, Dena Dubal, Shane Liddelow, Wenjie Luo, Li Gan

## Abstract

Demyelination occurs in aging and associated diseases, including Alzheimer’s disease. Several of these diseases exhibit sex differences in prevalence and severity. Biological sex primarily stems from sex chromosomes and gonads releasing sex hormones. To dissect mechanisms underlying sex differences in demyelination of aging brains, we constructed a transcriptomic atlas of cell type-specific responses to illustrate how sex chromosomes, gonads, and their interaction shape responses to demyelination. We found that sex-biased oligodendrocyte and microglial responses are driven by interaction of sex chromosomes and gonads prior to myelin loss. Post demyelination, sex chromosomes mainly guide microglial responses, while gonadal composition influences oligodendrocyte signaling. Significantly, ablation of the X-linked gene Toll-like receptor 7 (*Tlr7*), which exhibited sex-biased expression during demyelination, abolished the sex-biased responses and protected against demyelination.

**One-sentence summary:** Cell type-specific processes underlying aged demyelination are sex-biased and mediated by *Tlr7*.

## Introduction

Demyelination, or white matter loss, has been widely studied in autoimmune diseases such as multiple sclerosis (MS), which is much more common in women than in men. Emerging evidence has introduced the importance of demyelination in aging-associated neurodegenerative diseases, including Alzheimer’s disease (AD) and Parkinson’s disease (PD) (*1-6*). Like MS, AD and PD also exhibit striking sex differences. In AD, prevalence is also higher in women than in men, with two thirds of AD patients being women. In contrast, the opposite is true for PD with male patients predominant (*7-10*). Little is known about how biological sex shapes the mechanisms underlying demyelination with age.

Biological sex is primarily contributed by sex chromosomes and gonads, which release hormones such as estrogen and testosterone and are strongly influenced by aging. To investigate whether sex-based variations are influenced by sex chromosomes, gonads, or their combined interaction, we utilized the Four Core Genotype (FCG) mouse model, which allows for independent assessment of gonadal and sex chromosomal effects (*11*). In this model, the *Sry* gene required for testes formation is removed from the Y chromosome and inserted into an autosome, resulting in 4 sex chromosome-gonad combinations: XX with ovaries (XXO), XY with ovaries (XYO), XX with testes (XXT), or XY with testes (XYT). For the purposes of this study, XXO refers to female sex, and XYT male sex. Previous studies have shown that FCG mice with the same gonads exhibit similar levels of estrogen/testosterone regardless of sex chromosome background (*12-14*), allowing the assessment of sex chromosome effects (XX vs. XY) in the presence of similar levels of sex hormones, as well as gonadal effects (ovaries vs. testes). One source of sex-biased expression comes from incomplete X chromosome inactivation (XCI), which impacts at least 23% of X-linked genes (*15*). The resulting differences in gene expression between sexes could lead to skewing of disease mechanisms, resulting in differential outcomes (*16*). A catalogue of X-linked genes that exhibit sex-biased expression in various brain cell types during aged demyelination would be a valuable resource to dissect the contribution of sex chromosomes.

To induce demyelination and avoid autoimmune manifestations, we implemented cuprizone (CPZ) (*17*). Through single nuclei RNA sequencing (snRNAseq), spatial transcriptomics, and multiplex cytokine evaluations, we systematically studied CPZ-induced responses at 3 weeks (pre-demyelination, before myelin loss) and 5 weeks (post-demyelination, after myelin loss) specific to neural cell types, thereby pinpointing pivotal molecular mediators of sex-biased effects. Our comprehensive analyses showcase how sex chromosomes and gonads influence the central nervous system’s reactions to aged demyelination in a cell type-specific manner. We observed striking sex-biased responses in oligodendrocytes and microglia, which are key responders to demyelination and exhibit intrinsic sex differences (*18, 19*). Moreover, our study unveiled sex-biased expression of the X-linked microglial gene, *Tlr7*, during demyelination *in vivo* and in cultured microglia. Ablation of *Tlr7* abolished the sex differences in microglial and oligodendrocyte responses and protected against demyelination along with motor impairment, establishing an essential, detrimental role of *Tlr7* and its importance in dictating sex disparities in demyelination. This research paves the way for tailored therapeutics targeting sex differences, and identifying therapeutic targets that are applicable for both sexes.

## Results

### Sex-Biased Transcriptomic Changes Precede Myelin Loss in Aged Mice

CPZ-induced myelin loss is time-dependent; 3 weeks of exposure results in oligodendrocyte stress before myelin loss, and exposure to CPZ for 5 weeks or longer leads to severe myelin loss (*17*). To determine the influence of sex chromosomes or gonads in early-stage demyelination responses of aging mice, we treated 16–17-month-old FCG mice with 0.2% CPZ or control diet for 3 weeks (Fig. 1A-B) (*17*). As expected, we observed similar levels of estradiol in XXO and XYO mice (Fig. 1C), allowing the assessment of sex chromosomes in the same gonadal environment. Western blotting for myelin basic protein (MBP) in frontal cortex lysate revealed no significant differences induced by the CPZ diet after 3 weeks (Fig. 1D-E), consistent with previous studies (*20*). Immunostaining with MBP antibody also revealed no significant difference in myelin density across the hippocampal CA3 region and corpus collosum (Figure S1A-D).

**Figure 1:**
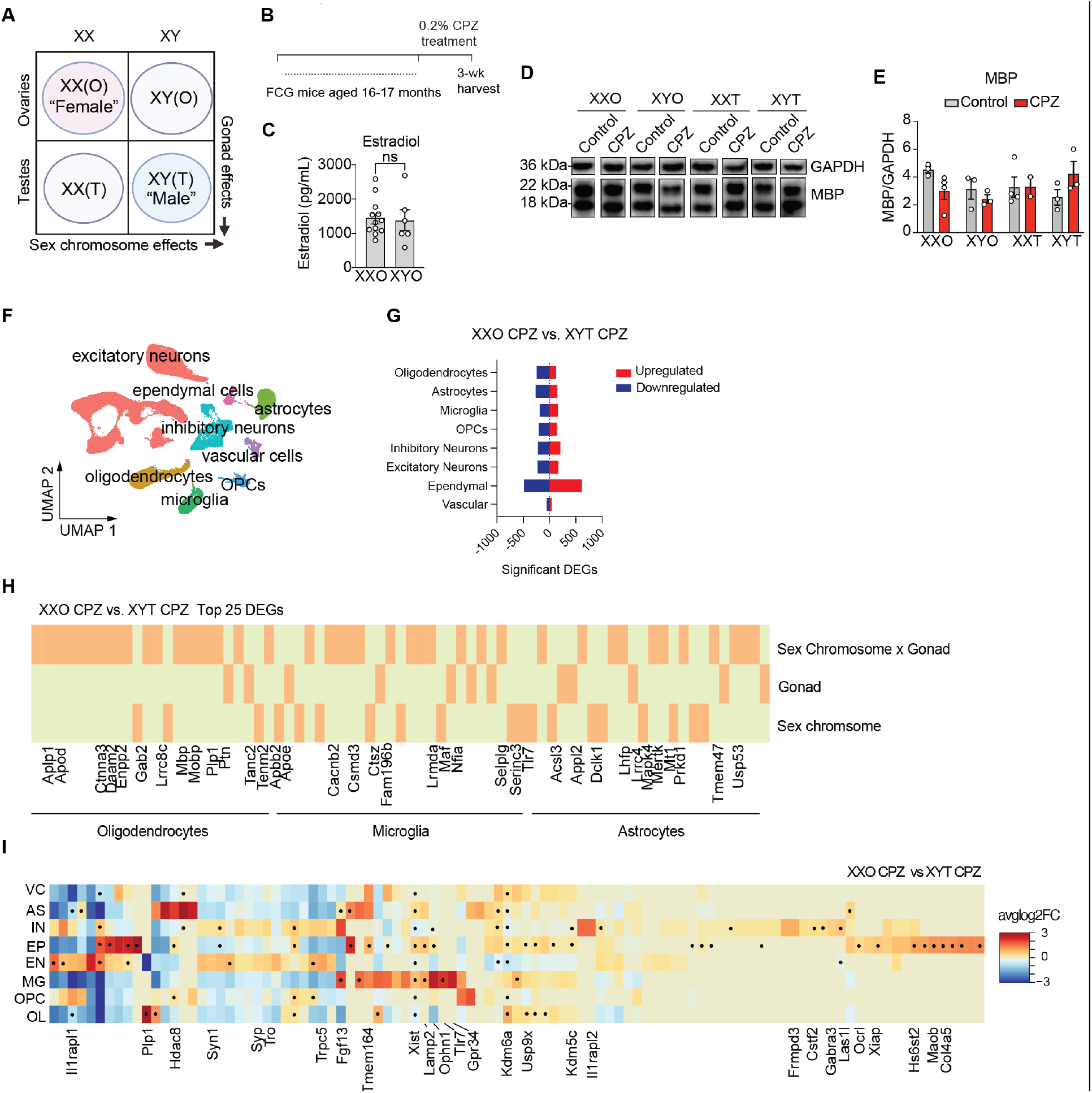
Sex-biased gene expression occurs in a cell type-specific manner pre-demyelination. (A) Schematic illustrating the Four Core Genotype comparison paradigm highlighting sex chromosome versus gonad comparisons. (B) Experimental design schematic. (C) Plasma estradiol concentration measured by ELISA in mice with ovaries treated with control and cuprizone diet. (D) Representative lanes of Western blot of MBP (18, 22kDa) and GAPDH (36 kDa) levels in frontal cortex lysate of 16–17-month-old mice treated with control or cuprizone diet and (E) quantification of MBP signal normalized to GAPDH. (F) UMAP of 8 detected cell types using single nuclei RNA-sequencing of the hippocampi from 16–17-month-old Four Core Genotype mice treated with control or cuprizone diet for 3 weeks, n=3 mice in each condition/genotype except XXT control, n=2. (G) Number of significant DEGs between XXO females treated with cuprizone and XYT males treated with cuprizone. (H) Diagram of top 25 DEGs between XXO cuprizone-treated and XYT cuprizone-treated samples for oligodendrocytes, microglia, and astrocytes. Orange fill indicates whether a given gene is primarily regulated by gonad, sex chromosome, or requires both according to sex chromosome-gonad scores (see Supp 3H and Methods for methods of calculation). (I) Heatmap of X-linked genes detected by cell type. Color indicates average log2foldchange expression of a given cell type compared to all other cell types. Black dot indicates that a given gene is also a significant DEG via MAST in XXO cuprizone-treated vs. XYT cuprizone-treated samples. All data are represented as mean +/-s.e.m., two-way ANOVA with Tukey’s post hoc multiple comparisons test.

We performed snRNA-seq to investigate early responses to CPZ. A total of 201,175 nuclei passed QC (Figure S2). The major cell types detected included oligodendrocytes, microglia, astrocytes, and neurons (Fig. 1F). Among them, oligodendrocytes exhibited the highest number (>700) of differentially expressed genes (DEGs) (Figure S3A), in line with oligodendrocytes being the major target of CPZ. When comparing female (XXO) and male (XYT) mice treated with CPZ, many sex-biased DEGs emerged across the different cell types (Fig. 1G, Table S1). Among the top 25 DEGs between females and males treated with 3 weeks of CPZ, we calculated whether the fold change difference for each gene was primarily attributed to sex chromosomes, gonads, or their interaction (Fig. 1H, Figure S3B). Across the glial cells of the brain, namely oligodendrocytes, microglia, and astrocytes, most sex-biased gene expression was cooperatively driven by gonads and sex chromosomes. Importantly, myelination genes such as *Mbp, Plp1*, and *Mobp* were widely downregulated in females compared to males and their expression was driven by sex chromosome-gonad interactions (Fig. 1H, Table S1). On the other hand, DEGs involved in the immune response expressed across microglia and astrocytes were largely influenced by either sex chromosomes (*Tlr7, Serinc3, Selplg*) or gonadal composition (*Ctsz, Tmem47, Prkd1*). AD-associated genes also appeared to be strongly regulated by either sex chromosomes (*Apbb2*) or gonads (*Apoe, Appl2*) (Fig. 1H, Table S1).

To begin to understand the role of sex chromosomes, we next examined how CPZ affects over 80 X-linked genes in males and females (Fig. 1I, Table S1). Notably, many X-linked genes exhibited sex-biased expression patterns in a cell type-specific manner, including a subset of X-linked genes, *Syn1, Syp*, and *Trpc5*, associated with synaptogenesis and synaptic activity (Fig. 1I). The oligodendrocyte gene *Plp1* was strongly downregulated in females, as well as *Kdm6a*, which has been implicated in AD-related longevity (*16*) (Table S1). The immune genes *Tlr7, Lamp2*, and *Tmem164* as well as the cell signaling genes *Ophn1* and *Gpr34* were among the most abundantly expressed by microglia compared to other cell types, with *Tlr7* and *Lamp2* also exhibiting sex-biased expression (Fig. 1I). Thus, these results show that X-linked genes can have both cell type- and sex-biased expression patterns in response to CPZ before the loss of myelin.

### Oligodendrocyte responses regulated by sex chromosomes and gonads pre-demyelination

We next used pseudobulk analysis to identify responses occurring across all oligodendrocytes. CPZ stimulated a sex-biased transcriptomic response, with about 500 more DEGs than observed in untreated oligodendrocytes (Figure S4A, Table S1). Top pathways downregulated by females with CPZ included neuronal support functions as well as myelination. The top pathways upregulated in females in response to CPZ were metabolism and transcription (Figure S4B–C).

We next performed sub-clustering of oligodendrocytes in the snRNA seq dataset and identified 6 subclusters of oligodendrocytes, including homeostatic (cluster 1) and myelinating oligodendrocytes (MOLs) (Fig. 2A, Figure S4D, Table S2). 4 out of 6 subclusters had populations altered by CPZ (Fig. 2B). CPZ treatment significantly reduced the number of cells in cluster 1 in XXO, XXT, and XYO mice, but not in male XYT mice (Fig. 2B). Thus, the preservation of the homeostatic oligodendrocyte subcluster appears to require both XY chromosomes and testes (Fig. 2B). Cluster 4 was induced by CPZ in females alone (XXO); Clusters 5 and 6 were induced by specific sex chromosome-gonad interactions but not in female (XXO) or male (XYT) mice (Fig. 2B).

**Fig 2:**
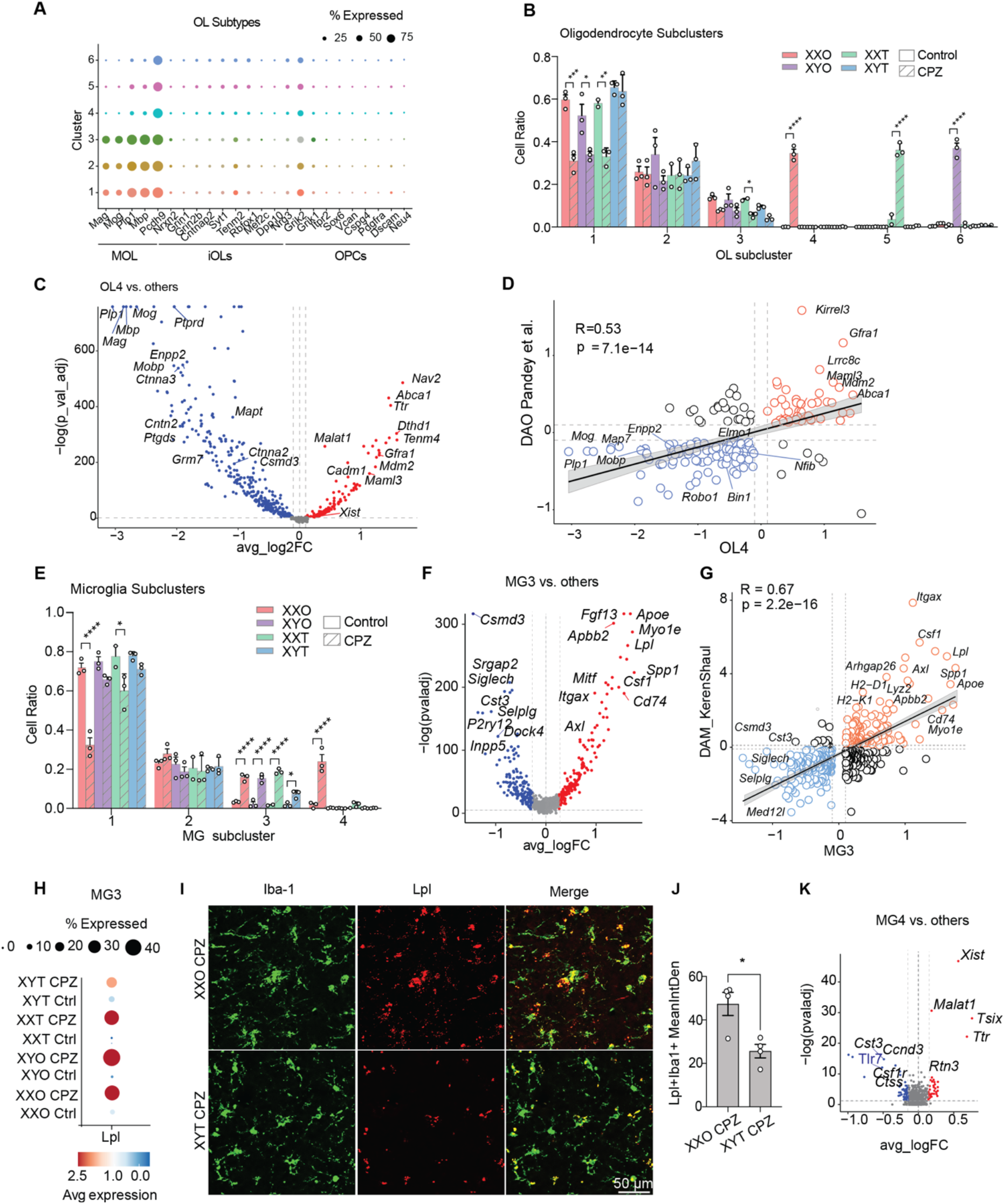
Sex chromosomes and gonads regulate oligodendrocyte and microglia responses pre-demyelination. (A) Dotplot of marker expression for myelinating oligodendrocytes, immature oligodendrocytes, and OPCs by subcluster. (B) Cell ratios of 30,011 oligodendrocytes within each subcluster colored by genotype and patterned by treatment. (C) Volcano plot of DEGs in cluster 4 compared to all other clusters. Dashed lines represent log2foldchange threshold of 0.1 and adjusted p value threshold of 0.05. (D) Correlation between Cluster 4 markers and published Disease Associated Oligodendrocyte genes from Pandey et al. 2022. Dashed lines represent log2foldchange threshold of 0.1. Red circles are genes upregulated by Cluster 4 and blue circles are genes downregulated by cluster 4. Pearson’s correlation test (two-sided). (E) Cell ratio of 12,473 microglia within each subcluster colored by genotype and patterned by treatment. (F) Volcano plot of DEGs in cluster 3 compared to all other clusters. Dashed lines represent log2foldchange threshold of 0.25 and adjusted p value threshold of 0.05. (G) Correlation between Cluster 3 markers and published Disease Associated Microglia genes from Keren-Shaul et al. 2017. Dashed lines represent log2foldchange threshold of 0.1. Red circles are genes upregulated by Cluster 4 and blue circles are genes downregulated by cluster 4. (H) Dotplot of *Lpl* expression by genotype and treatment within cluster 3 (I) Representative images of LPL and IBA-1 immunofluorescence taken in CA3 region of XXO and XYT cuprizone-treated samples. (J) Quantification of LPL and IBA-1 colocalization. (K) Volcano plot of DEGs in cluster 4 compared to all other clusters. Dashed lines represent log2foldchange threshold of 0.1 and adjusted p value threshold of 0.05. Pearson’s correlation test (two-sided). All data are represented as mean +/-s.e.m., for data in (B, E) significance was determined by two-way ANOVA with Tukey’s post hoc multiple comparisons test. In (J) significance was determined by unpaired, nonparametric student’s t-test with Mann-Whitney test, *p<0.05, **p<0.01, ***p<0.001, ****p<0.0001.

Examination of cluster 4 (OL4) revealed downregulation of myelinating genes, including *Mbp, Mag*, and *Plp1*. Myelin regulators such as catenins were also downregulated in this cluster (Fig. 2C, Table S2). In addition, genes involved in oligodendrocyte endocytosis, phospholipase D signaling, and cell adhesion, which are required for myelin morphogenesis, were diminished in cluster 4 (*21*) (Figure S4E). Strikingly, cluster 4 exhibited a strong, positive correlation with a disease-associated oligodendrocyte (DAO) signature identified in MS patients, including high levels of *Kirrel3* and low levels of *Robo1* (Fig. 2D) (*22*). Taken together, sex chromosomes and gonads work cooperatively to induce early sex-specific responses in male and female oligodendrocytes preceding demyelination.

### Microglial responses are regulated by sex chromosomes and gonads pre-demyelination

Pseudobulk analyses of microglia from the snRNA seq dataset revealed a sex-biased transcriptomic response with about 250 genes differentially expressed (Figure S4F). Pathway analyses of the DEGs revealed downregulated metabolic processes and upregulated phagocytosis and immune pathways (Figure S4G-H). Neuroimmune genes such as *Apoe, H2-K1, Axl, Myo1e, Trim14*, and *Cd9* were upregulated in females compared to males, indicating that microglia from female mice likely stimulated a stronger immune response to early demyelination before myelin loss (Table S1).

At the single-cell level, we identified 7 microglial subpopulations and observed significant changes in 3 clusters induced by CPZ (Fig. 2E, Figure S4I, Table S3). Interestingly, the proportion of homeostatic microglial cluster 1 was downregulated by CPZ only in genotypes with XX sex chromosomes, indicating the predominant influence of sex chromosomes over gonads in transforming microglial states away from the homeostatic state. The proportion of cells in cluster 3 was increased by CPZ across all genotypes but less induced in males (XYT). In cluster 3, we observed the upregulation of hallmark disease-associated microglia (DAM) genes such as *Apoe, Lpl*, and *Cd74* (Table S3). Cluster 3 DEGs indeed were positively correlated with the DAM gene signature initially observed in an amyloid mouse model (*23*) (Fig. 2F-G). Expression of *Lpl* was significantly elevated by CPZ in all genotypes except for XYT (male) mice (Fig. 2H). Immunostaining further confirmed that levels of LPL were higher in CPZ-treated female microglia compared with male counterparts (Fig. 2I-J).

Cluster 4, which was unique to XXO mice treated with CPZ, exhibited upregulated expression of epigenetic factors, such as *Xist* and *Tsix*, as well as downregulated expression of a specific subset of immune genes, including *Tlr7, Csf1r, Ctss*, and *Nfia*, compared to their expression in all other subclusters (Fig. 2K, Table S3). Together, these findings indicate that female oligodendrocytes and microglia exhibit larger loss of homeostatic and stronger gain of disease-associated transcriptomic profiles compared to male counterparts.

### Ovaries Worsen CPZ-induced Neuroinflammation and Myelin Loss

Our results thus far revealed that sex chromosomes and gonads interact to exert striking sex-biased transcriptomic responses induced by CPZ prior to myelin loss. To examine the effects of sex chromosomes and gonads in mice with myelin loss, we treated aged FCG mice with CPZ for 5 weeks. In addition to weight loss throughout the treatment, significant motor impairment was detected at the end of the 5-week treatment using rotarod test (Fig. 3A-B).

**Fig 3.**
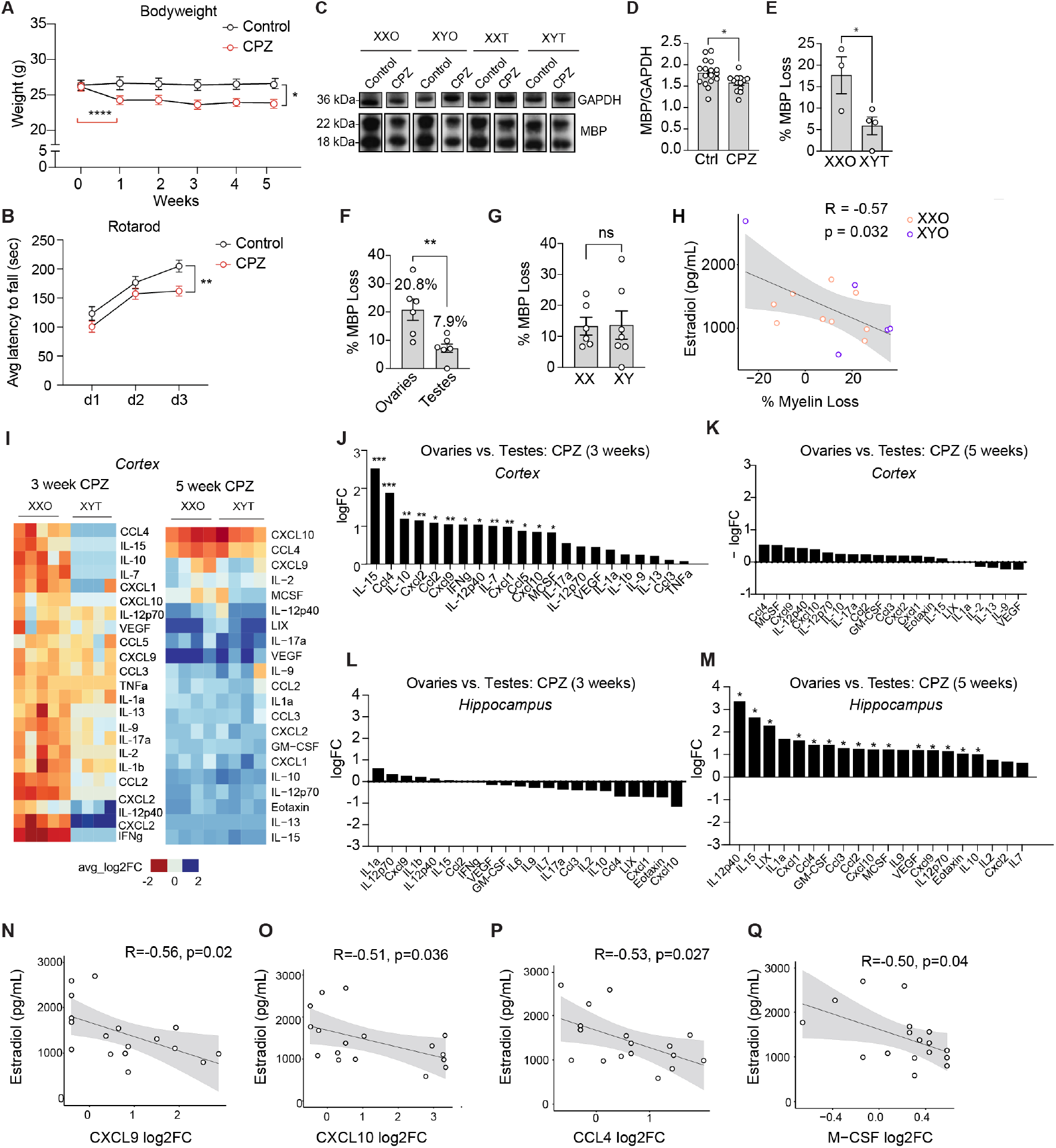
Ovaries elevated cytokines pre-demyelination and exacerbated myelin loss. (A) Weight change in control compared to 5-week cuprizone-treated FCG mice aged 10-11 months. (B) Average latency to fall during rotarod test for control and 5-week cuprizone-treated FCG mice, x-axis is day 1, day 2, day 3. (C) Representative lanes of Western blot for MBP (18 and 22 kDa) and GAPDH (36 kDa) performed on frontal cortex lysate of mice treated with control and 5-week cuprizone diet (n=3-6 mice per genotype per treatment). (D) Quantification of MBP signal between control and 5-week cuprizone-treated samples. (E-G) Quantification of percent myelin loss in 5-week cuprizone-treated samples from Western blot between females and males in (E), mice with ovaries versus testes in (F), and XX versus XY genotypes in (G). (H) Correlation between plasma estradiol concentration and percent myelin loss in 5-week cuprizone-treated mice with ovaries. (I) Heatmap of multiplex cytokine panel of frontal cortex lysate between cuprizone-treated XXO females and XYT males. (J) Quantification of cytokine levels average log2foldchange between genotypes with ovaries (XXO, XYO) and genotypes with testes (XXT, XYT) for 3 week and (K) 5-week CPZ treatments. (L) Quantification of hippocampal cytokine levels average log2foldchange between genotypes with ovaries (XXO, XYO) and genotypes with testes (XXT, XYT) for 3 week and (M) 5-week CPZ treatments. (N) Correlation of plasma estradiol level with levels of cytokine in hippocampal lysate CXCL9 (O) CXCL10 (P) CCL4 and (Q) M-CSF. Pearson’s correlation test (two-sided). All data are represented as mean +/-s.e.m. For data in (A-B) significance was determined by mixed model with Sidak’s post hoc multiple comparisons test. In (D-G) significance was determined by unpaired, nonparametric student’s t-test with Mann-Whitney test. Significance for cytokine levels was determined using linear regression package Glimma in R (Methods), *p<0.05, **p<0.01, ***p<0.001, ****p<0.0001.

In frontal cortex, CPZ-treated female mice showed significant myelin loss as detected by western blot, with female mice exhibiting greater myelin loss than males (Fig. 3C-E). Further analyses revealed that mice with ovaries (XXO and XYO) exhibited greater loss of myelin, suggesting a gonadal influence on demyelination (Fig. 3F). In contrast, the sex chromosome composition did not alter myelin loss (Fig. 3G). We measured plasma estradiol to determine whether systemic estradiol levels played a role in myelin loss in ovary-bearing mice. Interestingly, low estradiol was associated with more pronounced myelin loss in mice with ovaries (Fig. 3H).

To directly investigate the dynamic inflammatory responses during demyelination, we performed a multiplex ELISA assay to measure cytokine and chemokine levels 3 weeks and 5 weeks after CPZ treatment in both frontal cortex and hippocampus (Table S4). In frontal cortex, CPZ-treated female mice exhibited higher cytokine/chemokine levels than male equivalents at 3 weeks, but not 5 weeks, of CPZ treatment (Fig 3I). We also found that the presence of ovaries significantly increased several cytokines/chemokines after 3 weeks, but not 5 weeks, of CPZ treatment (Fig. 3J-K). In the hippocampus, however, ovary-associated neuroinflammation was minimal at 3 weeks of CPZ treatment but was profoundly higher at 5 weeks of CPZ treatment (Fig. 3L-M). Immunostaining of MBP revealed significant myelin loss in the hippocampus across all genotypes by the end of the 5 weeks (Figure S5A-B). Interestingly, we found that the levels of multiple cytokines in hippocampus, including CXCL9, CXCL10, CCL4, and M-CSF, were negatively correlated with plasma estradiol levels, indicating that estradiol loss could lead to exacerbated hippocampal neuroinflammation in response to myelin loss (Fig. 3N-Q).

### Spatial Transcriptomics Indicates Sex-biased Transcriptomic Responses Post-demyelination

The disparate dynamics of CPZ-induced demyelination in hippocampus and cortex prompted us to assess how sex chromosomes and gonads affect transcriptomic responses to myelin loss in a region-specific manner. We performed spatial transcriptomics in FCG mice treated with CPZ for 5 weeks, and sequenced coronal brain sections from XXO Ctrl, XXO CPZ, XYT Ctrl, and XYT CPZ animals (Figure S6). A total of 25 transcriptomic clusters were identified (Fig. 4A, Table S5). Based on the marker gene expression, several regions were identified, including white matter, hippocampus, and dorsal/ventral cortices (Fig. 4B). For the white matter cluster 7, comparing female and male mice treated with CPZ, we observed the downregulation of numerous ribosomal proteins as well as the upregulation of the AD risk genes *Apod* and *Bin1* in female mice (Fig. 4C). To determine the cell type composition of cluster 7, we cross-referenced cluster 7 marker genes with published cell type-specific signatures, including signatures for DAM, DAOs, disease-associated astrocytes (DAAs), and remyelinating oligodendrocytes (RemyeOL) (*22-25*) (Fig. 4D). Most overlapping genes in cluster 7 were DAM-related genes, but DAO- and RemyeOL-related genes were also represented, with few genes related to DAAs. When gene module scores for these signatures were mapped to spatial locations, we observed that DAMs were distributed throughout the brain, whereas DAOs were highly concentrated in the dorsal cortex and RemyeOLs were concentrated in white matter (Fig. 4D).

**Fig 4:**
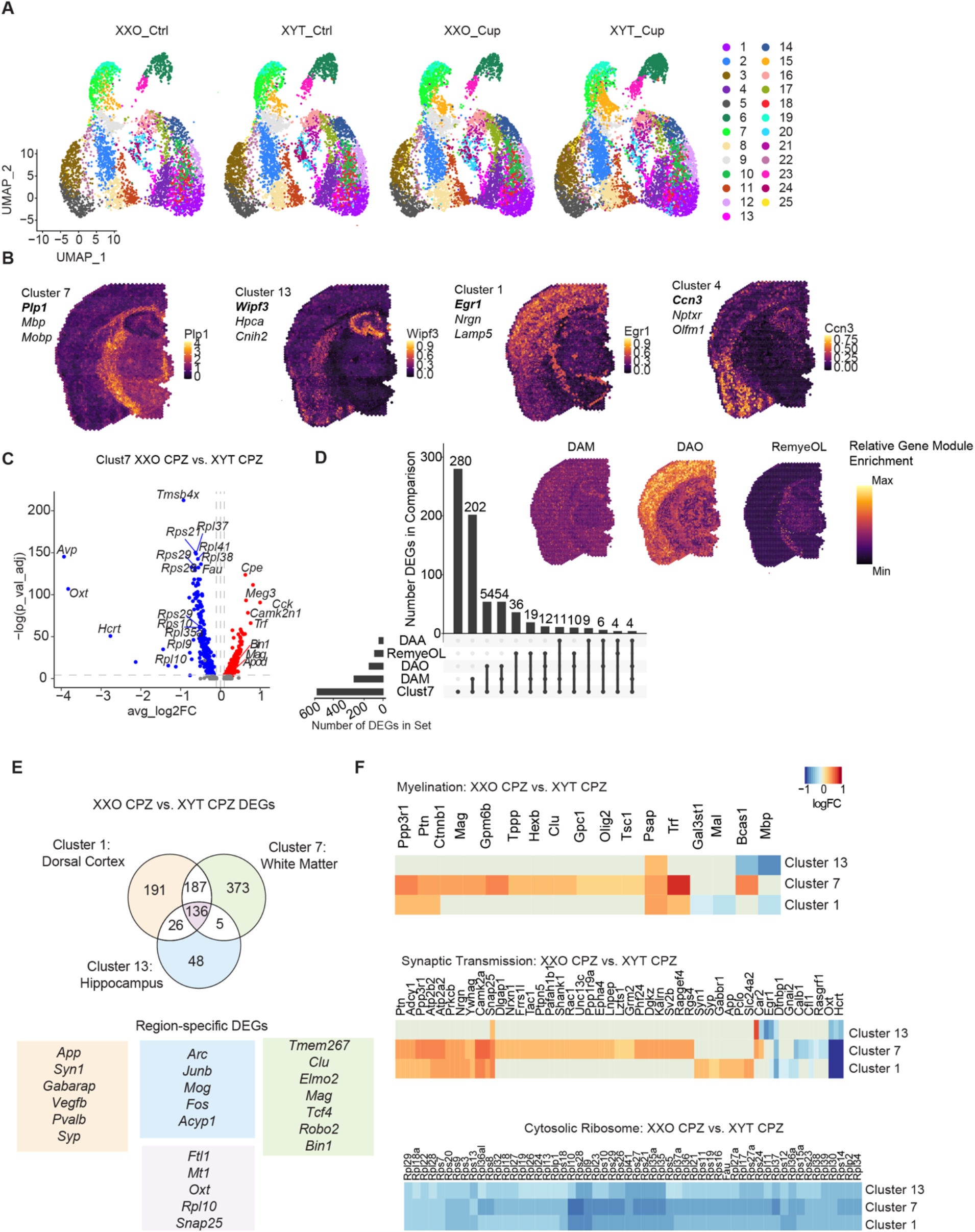
White matter, hippocampal, and cortical clusters exhibit sex-biased profiles post-demyelination. (A) UMAP of spatial subclusters across XXO control, XXO CPZ, XYT control, and XYT CPZ-treated 9–11-month-old FCG mice. (B) Representative section showing marker gene expression for clusters 7, 13, 1, and 4. Marker gene shown is bolded. (C) Volcano plot showing cluster 7 DEGs between XXO and XYT treated with 5-week cuprizone. (D) Upset plot showing cluster 7 overlap with published gene signatures for DAM, DAO, DAA, and RemyeOL. Representative section showing module enrichment scores for DAM, DAO, and RemyeOL are shown. (E) Venn diagram showing number of DEGs between females (XXO) and males (XYT) in clusters 1, 7, 13, and overlap. Unique DEGs for each region between XXO CPZ and XYT CPZ are shown with rectangle color corresponding to Venn diagram color. (F) Heatmaps for myelination, synaptic transmission, and cytosolic ribosome gene expression in clusters 1, 7, and 13.

To better understand whether sex-biased DEGs were region-dependent, we compared DEGs in white matter, hippocampus, and dorsal cortex clusters from female and male mice treated with CPZ (Table S5). White matter exhibited the most sexually dimorphic transcriptomic profile, followed by the dorsal cortex (Fig. 4E). Unique sex-biased genes in the dorsal cortex included *App, Syp*, and *Pvalb*, potentially indicating differences in synaptic transmission. Among the unique hippocampal genes were plasticity-oriented genes, such as *Arc* and *Fos*. The 3 regions also shared 136 sex-biased genes, including *Oxt, Snap25*, and *Mt1* (Table S5). Heatmaps show that genes related to myelination and synaptic transmission signaling were upregulated in females, particularly in white matter cluster 7 (Fig. 4F). Cytosolic ribosome genes were downregulated in females across all 3 regions (Fig. 4F).

### Sex Regulates Oligodendrocyte and Microglia Signaling During Demyelination

Our results from the spatial transcriptomics experiment revealed that sexual dimorphisms in transcriptomic profile occur across key brain regions during demyelination. Since oligodendrocytes are the primary glial cells that produce and maintain myelin sheaths, we examined oligodendrocyte responses at the single cell level at different stages of demyelination in hippocampus (Figure S7). We compared the snRNA seq datasets from 3-week CPZ and 5-week CPZ treatments (Table S6). Like the dynamic nature of inflammatory responses in cytokine levels, the oligodendrocyte transcriptome was also regulated in a temporal-specific manner. The difference between male and female oligodendrocytes was greatest after 3 weeks of CPZ treatment before myelin loss and lessened by 5 weeks of CPZ treatment (Fig. 5A). Nevertheless, we detected 49 genes, including the key myelination genes *Plp1* and *Mobp*, that exhibited sex-biased expression under both treatment conditions (Fig. 5A-B, Table S6). Interestingly, the sex-biased myelination genes were first downregulated in females following 3 weeks of CPZ treatment but were upregulated in females after 5 weeks of treatment, suggesting sex-specific regulation of these myelination genes at different demyelination stages (Fig. 5A-B). To determine which sex-biased transcriptomic differences were attributed to sex chromosomes versus gonadal composition, we compared mice with XX and XY genotypes and mice with ovaries and testes during demyelination (Table S6). Among the DEGs, 13 of the sex biased DEGs were influenced by sex chromosome composition, 60 were influenced by gonadal composition, and 20 were influenced by both (Fig. 5C). The 60 DEGs influenced by gonadal composition included myelination genes (*Mobp, Plp1*), amyloid genes (*Aplp1, Abca2, Itm2b*), and AD risk genes (*Apoe, Apod*) (Fig. 5C).

**Fig 5:**
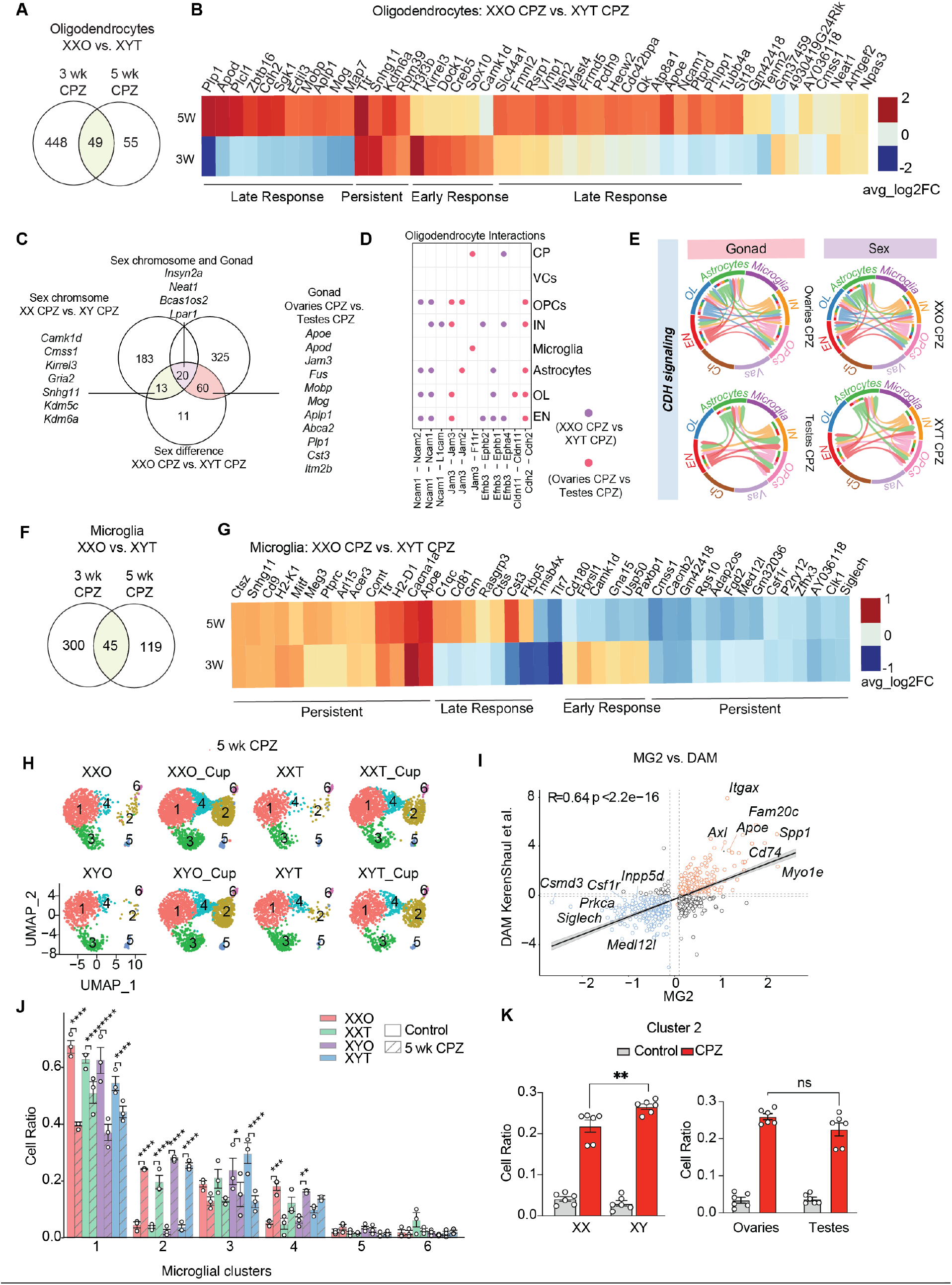
Regulation of oligodendrocytes and microglia by sex during demyelination. (A) Venn diagram showing number of oligodendrocytes DEGs between females (XXO) and males (XYT) in 3-week cuprizone-treated mice, 5-week cuprizone treated mice, and overlap. (B) Heatmap showing log2foldchange values of DEGs between females (XXO) and males (XYT) overlapping between 3- and 5-week cuprizone treatment in oligodendrocytes (overlap in Venn Diagram from A). (C) Venn diagram of number of DEGs between female and male, and DEGs contributed by sex chromosome (green), gonad (pink), and both (purple). (D) CellChat analysis showing ligand-receptor pairs with sex-biased expression (purple) or sex-biased and gonad-driven expression (pink). (E) Chord diagram showing CDH signaling split by sex, gonad, or sex chromosome. (F) Venn diagram showing number of microglia DEGs between females (XXO) and males (XYT) in 3-week cuprizone compared to control, 5-week cuprizone compared to control, and overlap. (G) Heatmap of DEGs between XXO females and XYT males treated with cuprizone overlapping between 3-week and 5-week cuprizone treated microglia. Heat indicates average log2foldchange expression of a given gene between XXO cuprizone-treated and XYT cuprizone-treated samples. (H) UMAP of 10,929 microglia split by condition (I) Correlation between Cluster 2 markers and published Disease Associated Microglia genes from Keren-Shaul et al. 2017. Dashed lines represent log2foldchange threshold of 0.1. Red circles are genes upregulated by Cluster 2 and blue circles are genes downregulated by cluster 2. (J) Cell ratios within each subcluster by genotype and treatment. (K) Cell ratios of cluster 2 in control and 5-week cuprizone-treated samples between XX genotypes (XXO, XXT) and XY genotypes (XYO, XYT). Adjacent graph shows cell ratios of cluster 2 between ovaries (XXO, XYO) and testes (XXT, XYT) genotypes. All data are represented as mean +/-range, for data in (J) significance was determined by two-way ANOVA with Tukey’s post hoc multiple comparisons test. For data in (K) significance was determined by unpaired, nonparametric student’s t-test with Mann-Whitney test, *p<0.05, **p<0.01, ***p<0.001, ****p<0.0001.

We next investigated how the sex-biased DEGs identified in oligodendrocytes influence other cell types using CellChat analysis, which provides predictions on cell-cell interactions (*26*). We focused analyses on sex biased DEGs and DEGs caused by gonadal composition, the two most abundant categories (Fig. 5D). Ligands whose expression was specifically driven by gonads included *Cdh2*, which has been shown to facilitate oligodendrocyte myelination capacity via neurons (*27*). Strikingly, the sex difference in CDH signaling appeared to be exclusively attributed to gonadal composition. CDH signaling was completely absent in male (XYT) oligodendrocytes and in the presence of testes. In female (XXO) oligodendrocytes, CDH signaling projected to astrocytes and neurons (Fig. 5E).

Microglia exhibited a robust sex-biased transcriptomic signature after myelin loss (Fig. 5F-G, Table S7). Neuroimmune genes such as *Apoe* and *H2-D1* remained highly upregulated in females compared to males, and the homeostatic microglial genes *P2ry12* and *Csf1r* were downregulated in females. The expression of smaller subsets of genes was exclusively observed pre- or post-demyelination; among these genes were *C1qc* and *Cd180* pre-demyelination along with *Usp50* post-demyelination.

At the single-cell level, we identified a subcluster of microglia with increased abundance induced by CPZ treatment across all genotypes, MG2, which exhibited a transcriptomic profile highly correlated to that of DAMs (Fig. 5H-J, Table S7). This cluster was sex chromosome-driven, but the number of cells within this cluster was decreased in mice with female XX sex chromosome composition (Fig. 5K). This cluster was not affected by gonadal composition (Fig. 5K). Within cluster 2, XX microglia exhibited differential expression of autosomal DAM-related genes in addition to X-linked genes such as *Tlr7, Kdm6a*, and *Tmsb4x* (Table S7). Taken together, these findings suggest that sex chromosomes appear to play a more important role than gonads in modulating microglial activation states than post-demyelination.

### Tlr7 KO Mitigates Transcriptomic Sex Differences and Protects Against Myelin Loss During Aging

During demyelination, myelin debris falling from the axon sheath stimulate microglia. To further dissect the contribution of sex chromosomes to microglial responses to demyelination, we treated primary male or female microglia from FCG mice with myelin. We performed bulk RNA-seq and compared transcriptomic responses in primary microglia isolated from female and male mice (Table S8). To explore converging mechanisms *in vitro* and *in vivo*, we analyzed the overlapping DEGs by comparing the DEGs across our 3 experimental conditions (*in vitro* myelin treatment, *in vivo* CPZ treatment for 3 weeks, and *in vivo* CPZ treatment for 5 weeks) (Fig. 6A, Table S8). Of these 17 DEGs, *Tlr7*, an X-linked immune gene that is predominantly expressed by microglia, was consistently and substantially downregulated in all 3 experimental groups (Fig. 6B).

**Fig 6.**
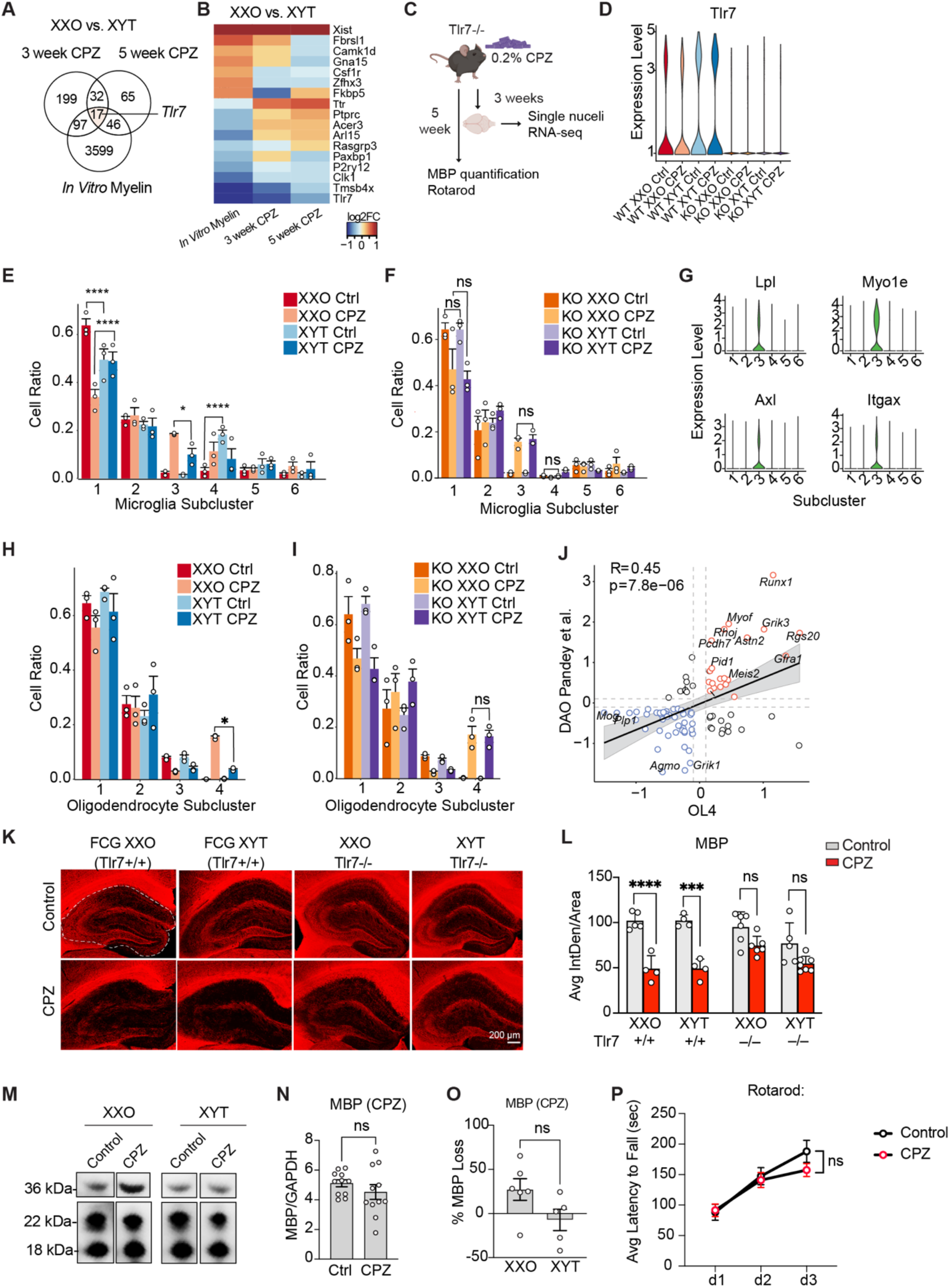
TLR7 KO abolished sex differences in pre-demyelination transcriptomic responses and protected against myelin loss. (A) Venn diagram of DEGs between XXO and XYT microglia treated with myelin *in vitro* or cuprizone *in vivo*. (B) Heatmap of 17 DEGs between XXO and XYT shared across *in vitro* myelin and *in vivo* cuprizone treatments. (C) Experimental design for cuprizone treatment of TLR7KO mice. (D) Violin plot showing pseudo-bulk *Tlr7* expression (E) Microglia cell ratios of each subcluster by genotype and treatment for Four Core Genotype samples treated with control or 3-week cuprizone diet. (F) Microglia cell ratios of each subcluster by genotype and treatment for *Tlr7* knockout samples treated with control or 3 week cuprizone diet. (G) Violin plots showing expression levels of DAM genes *Lpl, Myo1e, Axl*, and *Itgax* by subcluster. (H) Oligodendrocyte cell ratios of each subcluster by genotype and treatment for Four Core Genotype samples treated with control or 3-week cuprizone diet. (I) Oligodendrocyte cell ratios of each subcluster by genotype and treatment for *Tlr7* knockout samples treated with control or 3 week cuprizone diet. (J) Correlation between Cluster 4 markers and published Disease Associated Oligodendrocyte genes from Pandey et al. 2022. Dashed lines represent log2foldchange threshold of 0.1. Red circles are genes upregulated by Cluster 4 and blue circles are genes downregulated by cluster 4. (K) Representative images of MBP immunofluorescence in whole hippocampus of control and cuprizone-treated mice. (L) Quantification of MBP signal intensity. (M) Representative lanes of Western blot for MBP (18 and 22 kDa) and GAPDH (36 kDa) performed on frontal cortex lysate of mice treated with control and 5-week cuprizone diet (n=3-6 mice per genotype per treatment). (N) Quantification of MBP signal between control and 5-week cuprizone-treated samples. (O) Quantification of percent myelin loss in 5-week cuprizone-treated samples from Western blot between females and males in G. (P) Average latency to fall during rotarod test for control and 5-week cuprizone-treated mice, n=11-16 per treatment. All data are represented as mean +/-range, for data in (E-F, H-I) significance was determined by two-way ANOVA with Tukey’s post hoc multiple comparisons test. For data in (L), significance was determined by three-way ANOVA with Tukey’s post hoc multiple comparisons test. For data in (N-O) significance was determined by unpaired, nonparametric student’s t-test with Mann-Whitney test. For data in (P), significance was determined by mixed model with Sidak’s multiple comparisons test, *p<0.05, **p<0.01, ***p<0.001, ****p<0.0001.

Given the importance of *Tlr7* in microglial responses and the sex-biased inflammation during demyelination, we next tested if *Tlr7* contributed to the sex differences seen during demyelination. To determine if deleting *Tlr7* abolishes the sex differences in demyelination, we treated TLR7 knockout mice (TLR7KO) with CPZ for 3 or 5 weeks then assessed rotarod performance, myelin loss, and cell type-specific responses via snRNA-seq (Fig. 6C). The snRNA-seq data from male and female TLRKO mice treated with 3-wk CPZ were directly compared to that of FCG mice with intact *Tlr7*. For direct comparison, the two sets of snRNA-seq data were integrated; This integrated dataset contained 188,720 nuclei that passed QC (Figure S8). As expected, *Tlr7* RNA was not detected in the KO mice (Fig. 6D).

The integrated microglia population exhibited 6 subclusters (Figure S9A, Table S8). We observed a robust loss of the homeostatic microglia cluster (cluster 1) and an increase in DAMs-enriched cluster (cluster 3) in female mice but not in male mice treated with 3-week CPZ, reaffirming the sex-biased microglial response (Fig. 6E). In contrast, TLR7KO male and female microglial responses were similar, exhibiting similar trends toward decreases in cluster 1 and similar increases in cluster 3 (Fig. 6F). Analyses of cluster 3 revealed enrichment of typical DAMs, including *Lpl, Myeo1e, Axl*, and *Itgx* (Fig. 6G).

The integrated oligodendrocytes population exhibited 4 main subclusters (Figure S9B, Table S8); CPZ expanded the proportion of cells in cluster 4 in female, but not in male FCG mice with normal *Tlr7* as expected (Fig. 6H). The sex difference was abolished in the absence of *Tlr7*, as both males and female TLR7KO mice exhibited similar expansion of oligodendrocytes cluster 4 in response to CPZ (Fig. 6I). Cluster 4 is enriched with DAO, as expected (Fig. 6J, Figure S9C).

While TLRKO abolished the sex-dimorphic transcriptomic responses, resulting in similar responses in male and female microglia and oligodendrocytes at the early phase of demyelination, similar weight loss was induced by 5 weeks of CPZ regardless of the presence of *Tlr7* (Figure S8D). However, deleting *Tlr7* ameliorated CPZ-induced myelin loss in frontal cortex and hippocampus (Fig. 6K-O). Moreover, deleting *Tlr7* prevented motor impairment across males and females (Fig. 6P). Taken together, our results established the importance of *Tlr7* in mediating sex differences in demyelination and its essential role in contributing to demyelination and associated motor deficits.

## Discussion

Our current study combines snRNAseq, spatial transcriptomics, and behavioral studies to shed light on the roles of sex chromosomes and gonads in cell type-specific responses to disease progression related to demyelination in aging brains. We established the first brain cell type-specific atlas that allows for dissociating the impact of sex chromosomes and gonads on demyelination, identifying notable roles of sex chromosomes and gonadal composition in influencing the responses of oligodendrocytes and microglia. A key discovery was the sex-biased expression of the microglial gene, *Tlr7*. Removing TLR7 eradicated the observed sex differences in cell type-specific responses, offering protection against demyelination and motor impairment.

A prior study documented the sex-specific expression of X-linked genes in various human tissues (*15*). Since sex-specific gene expression is tied to varying susceptibilities to immune diseases, especially autoimmune conditions (*28*), our current study mapped the expression of over 80 X-linked genes in different brain cell types at the single cell level, both under standard conditions and after demyelination events in aging. We found that when faced with demyelinating triggers, sex-specific gene expression spans disease-associated processes including myelin formation (*Plp*), chaperon-mediated autophagy (*Lamp2*), and inflammatory reactions (*Tlr7*). These revelations present opportunities to further investigate the molecular underpinnings of sex-specific neuroinflammatory and demyelinating events. The regulation of XCI is a nuanced process, influenced by the interplay of non-coding RNAs, epigenetic alterations, and nuclear structure, some of which could be involved in meditating sex-specific neuroinflammation and demyelination.

Our study constructed a dynamic cell type-specific map of responses shaped by sex chromosomes and gonads during the demyelination process. Striking sex-biased responses were observed in both microglia and oligodendrocytes prior to overt myelin loss, with female oligodendrocytes and microglia exhibiting robust expansion of disease-associated states while little shifts were observed for male counterparts. These findings are consistent with the notion that the disease-associated transcriptomic changes in microglia and oligodendrocytes are not only responses to the injury, but also active contributors to disease progression. Indeed, the stronger transcriptomic responses in female mice are associated with more significant myelin loss than male mice. Interestingly, while both sex chromosomes and gonads contribute to the robust transcriptomic responses in females, gonads, especially ovaries, play a pivotal role in demyelination during aging. In mice with ovaries (XXO and XYO), levels of plasma estradiol, a type of estrogen, seem to inversely impact myelin loss. Moreover, levels of specific cytokines were inversely correlated with plasma estradiol levels at the timepoint with significant myelin loss in the hippocampus, indicating that a reduction in estradiol may intensify hippocampal neuroinflammation in response to myelin loss. These observations align with the abundant evidence that estrogen is neuroprotective (*29*). Specifically, estrogen receptor ERß on the microglia and oligodendroglia were found to contribute to the neuroprotective/anti-inflammatory and remyelinating effects of estrogen in female mice (*30*). However, it is also conceivable that other gonadal hormones, such as progesterone, testosterone, follicle-stimulating hormone (FSH), or luteinizing hormone, may also contribute the differences in myelin loss attributed to ovaries. For example, a recent study showed a combined beneficial effect of testosterone and estrogens on microglial responses favoring regeneration in a mouse model of EAE (*31*). Reduced estrogen can also induce higher levels of FSH, which were found to impair cognition and exacerbate AD pathology in an ovariectomized mouse model of AD (*32*).

Sex-biased microglial and inflammatory responses to CPZ were also dynamic and region-specific. In frontal cortex, heightened cytokine/chemokine levels, many of which are pro-inflammatory, were associated with ovaries after 3 weeks (but not 5 weeks) of CPZ treatment. Contrastingly, in the hippocampus, ovary-related neuroinflammation was almost negligible at 3 weeks but notably high after 5 weeks. Moreover, levels of specific cytokines inversely correlated with plasma estradiol levels when there is significant myelin loss in the hippocampus, consistent with anti-inflammatory function of estrogen. Interestingly, the strong DAM responses at 3 weeks of CPZ appear to be driven by the sex chromosomes and gonads interaction, while at 5 weeks of CPZ the microglial responses seemed to be driven by sex chromosomes, with lower responses associated with XX chromosomes. Dissecting how sex modifies the link between inflammatory responses and myelin loss will shed light on mechanisms underlying the dynamic demyelination process.

Our study uncovered TLR7 as a key player mediating the sex differences and myelin loss induced by CPZ. TLR7 recognizes fragments of single-stranded RNA (ssRNA) and triggers inflammatory responses (*33*). The ssRNA could come from RNA viral infection, such as SARS-COVID 2, or released from mitochondria (*33*). Our current study revealed that TLR7KO eliminated the sex differences in oligodendrocytes and microglia response to demyelination, suggesting that TLR7 plays an important role in the sex-biased demyelination response. Moreover, TLR7 rescued myelin loss and motor deficits in both males and females treated with CPZ, pointing to a key detrimental role of TLR7 during demyelination. *Tlr7* was found to facilitate TNFa-related pain resolution, which exhibit striking sex differences (*34*). Deleting *Tlr7* slowed down the adaptive resolution of pain in models of acute and chronic inflammation in both sexes (*34*), supporting a central role of TLR7 in determining the functional consequence of inflammation. How female microglia exhibit lower expression of TLR7 during demyelination is puzzling. One likely mechanism is sex-biased epigenetic regulations. In support of this hypothesis, a recent study showed that TLR7 has altered CpG methylation patterns, which leads to downregulation in males compared to females in the most severe cases of COVID-19 (*35*). Further studies are needed to determine whether epigenetic mechanisms are involved in sex-biased expression of TLR7 microglia. Intriguingly, miRNAs have emerged as endogenous ligands for *Tlr7* (*36*). It remains a possibility that the sex-biased expression of miRNAs in microglia plays a role in driving *Tlr7* activation during demyelination (*37*).

Sex differences in neuroscience remain vastly understudied, especially in the context of aging and disease. Our findings indicate that sex is a primary modifier of the demyelination process, which occurs in MS, AD, and PD, all of which exhibit sex-biased prevalence and severity (*38*). Thus, this study reinforces the importance of including biological sex as a variable in experimental designs, especially those pertaining to disease-associated models. The dynamic transcriptomic changes we observed in both oligodendrocytes and microglia highlight the nuanced differences in male and female responses. The fact that certain genes and cellular signaling pathways exhibit differential activity based on sex chromosomes or gonadal composition has profound implications for understanding disease mechanisms and developing potential treatments. Thus, these observations stress the need for consideration of biological sex in both basic and translational scientific studies.

## Supporting information

Materials/Methods, All Supplementary Figures

## ACKNOWLEDGEMENTS

The study was supported by:

the NIH U54NS100717 (to LG)

R01AG072758 (to LG)

R01AG054214 (to LG)

R01AG074541 (to LG)

Tau Consortium (to LG)

JPB Foundation (to LG)

Gilliam Fellowship (to CLL)

We would like to thank Fenghua Hu for assistance troubleshooting the MBP staining.

## AUTHOR CONTRIBUTIONS

Conceptualization: LG, CLL, LK

Methodology: PY, ERT, GAM, RDK, DD, SAL

Investigation: CLL, LK, LF, MYW, NRF, LJ, FY, JZ, and KN

Visualization: CLL, MYW, NRF

Funding Acquisition: LG, CLL

Writing – original draft: CLL, LG

Writing – reviewing and editing: LG, CLL, LK, SAL

## DECLARATION OF INTERESTS

The authors declare no competing interests.

## SUPPLEMENTARY MATERIALS

Materials and Methods

Figs. S1 to S9

Tables S1 to S8

References(*39-44*)

## Notes

### Competing Interest Statement

The authors have declared no competing interest.

