## Supplementary material for "Sex Chromosomes and Gonads Shape the Sex-Biased Transcriptomic Landscape in Tlr7-Mediated Demyelination During Aging": Materials/Methods, All Supplementary Figures

### Materials and Methods

*Mice*

Animal studies were carried out in the Research Animal Resources and Compliance facility. All experiments were performed in compliance with RARC standards for animal care. 3-month old Four Core Genotype males were generously donated by Dr. Dena Dubal from UCSF to establish the line by breeding XYT males with C57BL/6J females. B6.129S1-*Tlr7*$^{tm1Flv}$/J mice were purchased from Jackson Laboratory. Mice were maintained in barrier facility with 12-hour light, 12-hour dark cycle and stable temperature conditions.

Mice were euthanized using FatalPlus and perfused with cold PBS following cardiac puncture for blood collection. Blood was collected in EDTA-coated tubes and spun down to isolate plasma. After complete perfusion, left hemibrain was fixed in formalin for 24-48 hours followed by storage in 30% sucrose. Right hemibrain was microdissected for hippocampus and frontal cortex, which were frozen in dry ice and stored at -80°C until further processing.

*CPZ Treatment*

0.2% CPZ and batch-matched control chow was purchased from Teklad and stored at 4°C for <6 months. We requested that CPZ be mixed in with standard PicoLab 5053 already used in animal facility so mice were not exposed to new base chow. Diet was weighed and replaced every 2-3 days. Mice were weighed every week during total treatment time of 3-5 weeks.

*Single-nuclei RNA sequencing (snRNA-seq)*

Nuclei isolation from frozen mouse hippocampi was adapted from a previous study [Habib 2017], with modifications. All procedures were performed on ice or at 4°C. In brief, postmortem brain tissue was placed in 1,500 µl of Sigma nuclei PURE lysis buffer (Sigma, NUC201-1KT) and homogenized with a Dounce tissue grinder (Sigma, D8938-1SET) with 15 strokes with pestle A and 15 strokes with pestle B. The homogenized tissue was filtered through a 35-µm cell strainer, centrifuged at 600g for 5 min at 4 °C and washed three times with 1 ml of PBS containing 1% bovine serum albumin (BSA, Thermo Fisher Scientific, 37525), 20 mM DTT (Thermo Fisher Scientific, 426380500) and 0.2 U $\mu l^{-1}$ recombinant RNase inhibitor (Ambion, AM2684). Nuclei were then centrifuged at 600g for 5 min at 4 °C and resuspended in 350 µl of PBS containing 0.04% BSA and 1× DAPI, followed by fluorescence-activated cell sorting to remove cell debris. The sorted suspension of DAPI-stained nuclei was counted and diluted to a concentration of 1,000 nuclei per µl in PBS containing 0.04% BSA.

For droplet-based snRNA-seq, libraries were prepared with Chromium Single Cell 3′ Reagent Kits v3 (10x Genomics, PN-1000075) according to the manufacturer's protocol. The snRNA-seq libraries were sequenced on a NovaSeq 6000 sequencer (Illumina) with 100 cycles.

*snRNA-seq Data Processing*

Gene counts were obtained by aligning reads to the mm10 genome with Cell Ranger software (v.3.1.0; 10x Genomics). To account for unspliced nuclear transcripts, reads mapping to pre-mRNA were counted. Cell Ranger 3.1.0 default parameters were used to call cell barcodes. We further removed genes expressed in no more than three cells, cells with unique gene counts over 4,000 or less than 300, cells with UMI counts over 20,000 and cells with a high fraction of mitochondrial reads (>5%). Potential doublet cells were predicted using DoubletFinder for each sample separately, with high-confidence doublets removed. Normalization and clustering were done with the Seurat package v3.2.2 (*39*). In brief, counts for all nuclei were scaled by the total library size multiplied by a scale factor (10,000) and transformed to log space. A set of 2,000 highly variable genes was identified with SCTransform from the sctransform R package FindVariableFeatures function with vst method. This returned a corrected unique molecular identifier count matrix, a log-transformed data matrix and Pearson residuals from the regularized negative binomial regression model. Principal-component analysis was done on all genes, and t-distributed stochastic neighbor embedding was run on the top 15 principal components. Cell clusters were identified with the Seurat functions FindNeighbors (using the top 15 principal components) and FindClusters (resolution = 0.1). In this analysis, the neighborhood size parameter pK was estimated using the mean variance-normalized bimodality coefficient (BCmvn) approach, with 15 principal components used and pN set as 0.25 by default. Sample integration was performed using FindIntegrationAnchors and IntegrateData functions in Seurat. For the 3 week CPZ cohort snRNA-seq, we sequenced and integrated samples from XXO control (n=3), XXO CPZ (n=3), XXT control (n=2), XXT CPZ (n=3), XYO control (n=3), XYT control (n=3) and XYT CPZ (n=3). For the 5 week cohort, we sequenced and integrated samples from XXO control (n=3), XXO CPZ (n=3), XXT control (n=3), XXT CPZ (n=3), XYO control (n=3), XYT control (n=3) and XYT CPZ (n=3). For each cluster, we assigned a cell-type label using statistical enrichment for sets of marker genes and manual evaluation of gene expression for small sets of known marker genes. Differential gene expression analysis was done using the FindMarkers function and MAST (*40*). To identify gene ontology and pathways enriched in the DEGs, DEGS were analyzed using the MSigDB gene annotation database (*41, 42*). To control for multiple testing, we used the Benjamini–Hochberg approach to constrain the FDR.

*Spatial Transcriptomics*

Spatial transcriptomics was performed using the 10x Genomics' Visium platform following the manufacturer's instructions. Fresh frozen brain sections from Four Core Genotype males and females either treated with control or CPZ diet were cryosectioned coronally to 10 µm and mounted on the Visium Spatial Gene Expression Slide (n = 2-3 for each condition, 11 sections total). After methanol fixation at –20°C for 30 minutes, sections were stained with hematoxylin and eosin, imaged on a Keyence BZ-X710 using a 20x objective, and tiled using BZX710 software. Sections were then enzymatically permeabilized for 12 minutes, and spatial barcodes and unique molecular identifiers were added to the captured polyA mRNA. Libraries were generated using Visium Spatial Gene Expression Reagent Kits (10x Genomics, PN-1000184) with dual index primers (10x Genomics, PN-1000215) and sequenced on a NovaSeq6000 using a S4 100 cycle flow cell. The sequencing output was aligned to the mouse reference genome mm10 using SpaceRanger. The following libraries were used to analyze and visualize the data in R (4.1.0): Seurat (4.0.6), SingleCellExperiment (1.16.0), Harmony (0.1.1), BayesSpace (1.2.1), viridis (0.6.2), ggplot2 (3.3.5), purrr (0.3.4), scater (1.22.0), UpSetR (1.4.0), cowplot (1.1.1), ggpubr (0.4.0). SCTransform was applied to each sample separately before merging. For all downstream clustering analyses, variable features from all 11 samples were used. Harmony was used to resolve sample variability in 6 Harmony components and a resolution of 0.8 were used to cluster the spots across all samples. Seurat's FindMarkers function was used to identify differentially expressed genes in each cluster. BayesSpace (*43*) was used to identify sub-spot level gene expression. Seurat's AddModuleScore was used to visualize groups of genes in Visium brain sections.

*Primary Microglia Isolation and Culture*

Primary microglial cells were collected from mouse pups at postnatal days 1–3. Briefly, the brain cortices were isolated and minced. Tissues were dissociated in 0.25% Trypsin-EDTA for 10 min at 37 °C and agitated every 5 min. Two hundred microliters of DNAse I (Millipore) was then added. Trypsin was neutralized with complete medium (DMEM; Thermo Fisher) supplemented with 10% heat-inactivated FBS (Hyclone), and tissues were filtered through 70-µm cell strainers

(BD Falcon) and pelleted by centrifugation at 250*g*. Mixed glial cultures were maintained in growth medium at 37 °C and 5% $CO_2$ for 7–10 d *in vitro*. Once bright, round cells began to appear in the mixed glial cultures, recombinant mouse granulocyte–macrophage colony-stimulating factor (1 ng $ml^{-1}$; Life Technologies) was added to promote microglia proliferation. Primary microglial cells were collected by mechanical agitation after 48–72 h and plated on poly-D-lysine-coated 24-well plates (Corning) in growth medium. Microglia were maintained in phenol red-free DMEM supplemented with 10% FBS, 100 U $ml^{-1}$ penicillin and 100 μg $ml^{-1}$ streptomycin.

*Myelin Treatment*

Primary microglia were treated with 100 ug/mL myelin isolated from 2-3 month old WT C57/BL6J mice. After 24 hours, RNA was isolated (Zymo Research) and submitted to Novogene for analysis of RNA quality and integrity. All samples passed quality control, and RNA-seq libraries were prepared for sequencing using HiSeq.

*Western Blot*

25 ug frontal cortex or hippocampal lysate prepared above were boiled for 5 minutes and run on 26-well 4-12% Bis-Tris gels (Invitrogen) using MES buffer (Invitrogen) for 1 hour. Proteins were transferred from gel onto PVDF membrane for 2 hours. Membranes were washed three times for 10 min each in TBS with 0.01% triton X-100 (TBST). Membranes were blocked for 30 minutes in 5% milk in TBST and incubated with MBP (1:1,000, EMD Millipore), TLR7 (1:800, Novus Biologics), GAPDH (1:1,000, GeneTex) antibodies overnight in cold room. The following day, membranes were washed three times for 10 min each in TBST and incubated in appropriate HRP secondary for 1 hour, rinsed, and followed by ECL development and imaging using Bio-Rad imager.

*Immunofluorescence*

Fixed hemibrain was sectioned in 30 um-thick sections via microtome and stored in cryoprotectant. 3-5 hippocampal sections were picked per animal and following rinses with PBST, underwent antigen retrieval using Reveal Decloaker (Biocare Medical). Sections were incubated overnight at 4 degrees in primary antibodies: MBP (Millipore, 1:400), IBA-1 (Abcam, 1:400), LPL (Abcam, 1:100). Following Alexafluor secondary antibody incubation at 1:500, sections were rinsed with PBST and mounted using Prolong Gold Anti-Fade reagent with DAPI. Slides were sealed with clear nail polish and kept at 4°C.

*Image Analysis*

Immunofluorescence was visualized using LSM880 laser scanning confocal microscope and 3-5 images per animal were captured at 40X in CA3 of hippocampus and corpus collosum. Z stack of 3-5 planes was captured and for corpus collosum, 2 images were stitched to capture the region. Whole hippocampal images were taken at 10X with 11-13 planes and 2x2 images were stitched. After image capture, images were separated by color channel and underwent background subtraction and thresholding. Signal intensity was measured via ImageJ. Intensity of 1-2 control sections stained only with secondary antibody was averaged and subtracted from all sections. Signal intensity was normalized to area, averaged across sections for each animal, and plotted in Prism. 3D structure of microglia was reconstructed using Imaris software as previously described (*44*). Experimenters performing imaging and quantification were blinded.

*Pathway Analysis*

The EnrichR package was used to identify GO pathways corresponding to DEG lists generated by Seurat package. For larger gene lists (>200 genes), DEGs were entered into Gene Ontology database with threshold of log2FC <-0.1 or >0.1 and adjusted p val. < 0.05.

*CellChat Analysis*

The CellChat package was used to identify cell-cell interactions pathways corresponding to snRNA-seq data generated by Seurat package. Briefly, the QCed Seurat object containing all cell types and genes was converted into a CellChat object. Bubble chart was calculated based on significant ligand-receptor interactions being sent from oligodendrocytes only. The same package was used to generate chord diagrams of pathway activity by cell type.

*Rotarod Test*

During the final three days of CPZ treatment, mice underwent rotarod testing. After habituation for 1 hour, mice were placed on rotarod device accelerating to a maximum speed of 50 rpm. Mice were allowed to stay on device for a maximum of 5 minutes, with latency to fall recorded for each mouse. Mice completed three trials per day with at least 5 minutes between each trial. Three trials were repeated each day for a total of nine trials. Latency to fall was averaged per mouse and plotted.

*Multiplex ELISA*

Frontal cortex or hippocampal tissue was weighed and submerged in RIPA buffer containing protease inhibitors. Tissue was homogenized via sonication in 4 degree C water bath. BCA was performed to determine protein concentration. 50 ug protein per sample was used for Multiplex ELISA via Luminex MAGPIX mouse 32-plex kit. 23 cytokines were detected above low threshold values including: IL-15, Ccl4, IL-10, Cxcl2, Ccl2, Cxcl9, IFNg, IL-12p40, IL-12p70, Cxcl1, Ccl5, Cxcl10, MCSF, IL-17a, VEGF, IL-1a, IL-1b, IL-9, IL-13, Ccl3, TNFa, and IL-2.

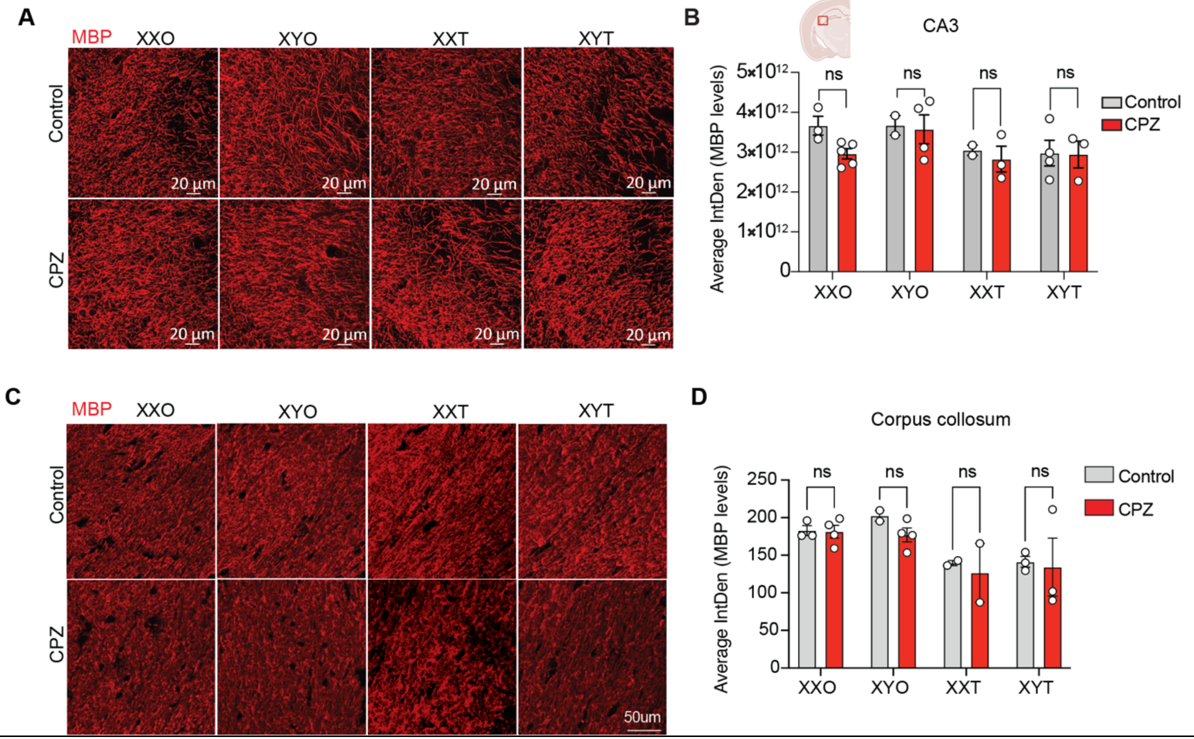


**Figure S1: 3-week cuprizone treatment does not result in myelin loss.** (A) Representative images of MBP immunofluorescence in hippocampal CA3 region of control and cuprizone treated mice. (B) Quantification of MBP signal intensity. (C) Representative images of MBP immunofluorescence in corpus collosum of control and cuprizone treated mice. (D) Quantification of MBP signal intensity. All data are represented as mean +/- s.e.m., two-way ANOVA with Tukey's post hoc multiple comparisons test.

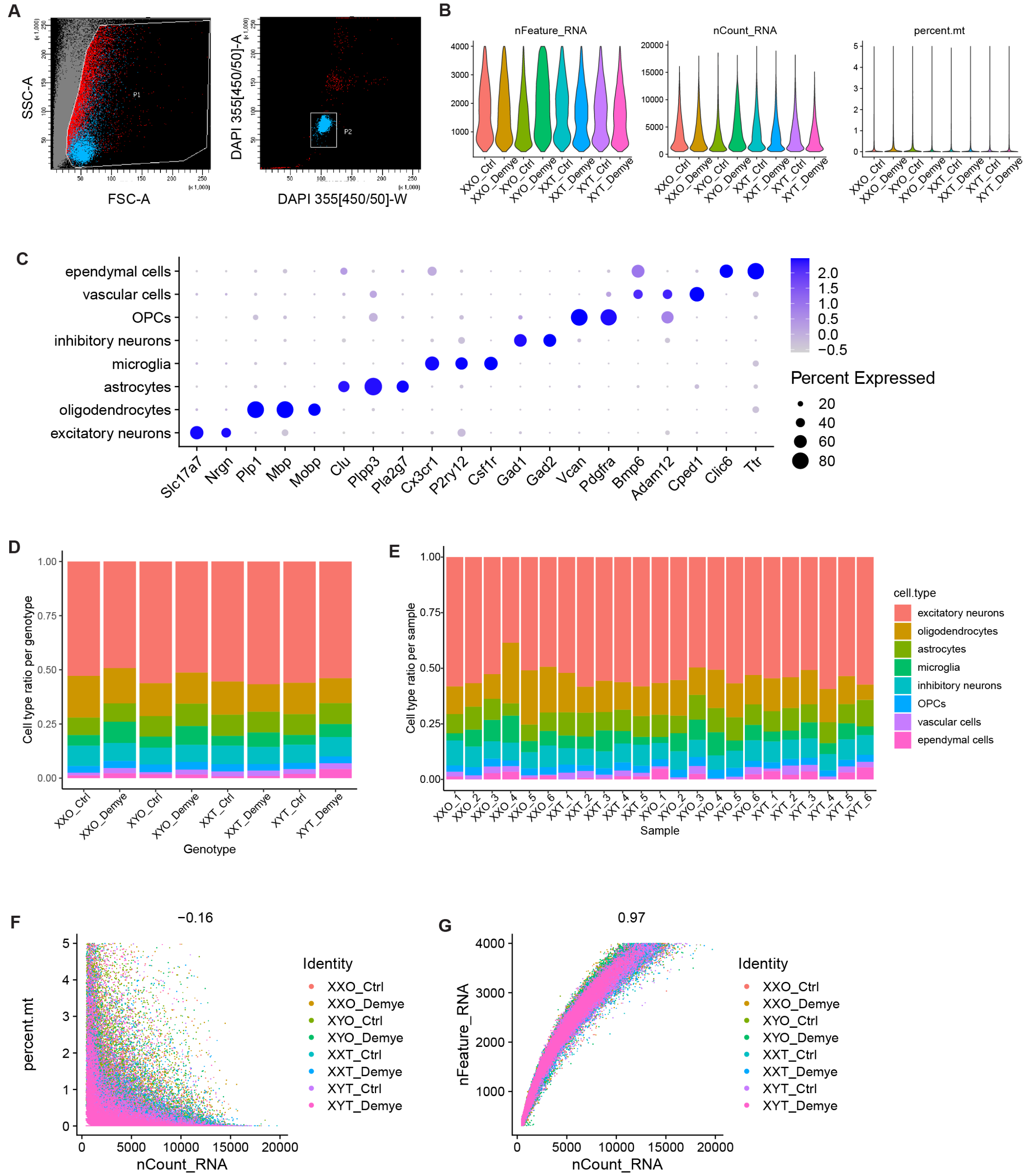


**Figure S2: Quality control assessment of single-nuclei RNA-seq of 3-week CPZ for FCG cohort.** (A) Representative FACS plot showing gating strategy used for collection of DAPI-labeled hippocampal nuclei after mechanical dounce homogenization. n=2-3 per genotype per treatment. (B) Quality-control plots showing equivalent amounts of total RNA features, total number of RNA counts, and percent mitochondrial RNA in nuclei used for downstream analyses. (C) Summary of genes used for cluster classification into different cell types. (D) Proportion of each cell type detected across the different genotypes. (E) Proportion of each cell type detected across the different samples. (F) Correlation between RNA count and percentage of mitochondrial genes per nuclei and (G) RNA count and RNA features for all samples.

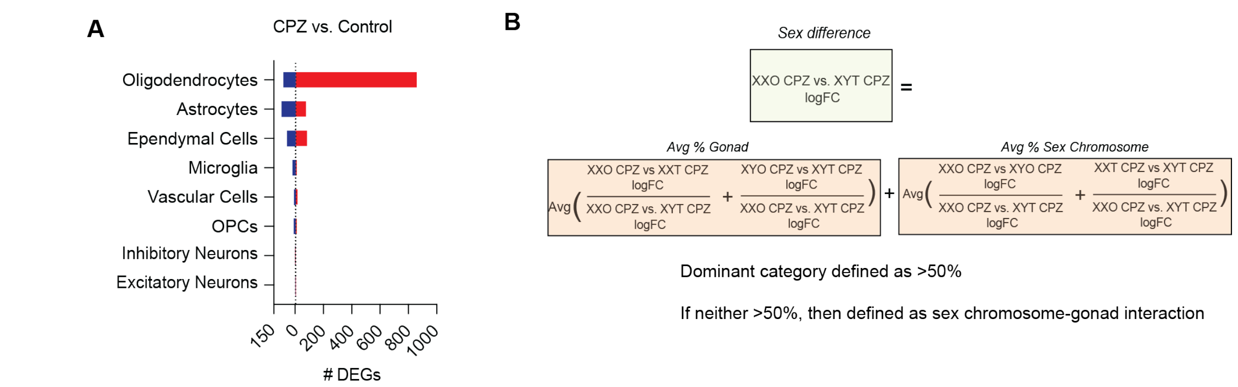


**Figure S3: Sex chromosomes and gonads modify transcriptomes of oligodendrocytes and microglia pre-demyelination.** (A) Number of significant DEGs by cell type in cuprizone (n=12) vs. control (n=12) treated samples with log2FC cutoff <-0.25 or >0.25. (B) Equation for calculating gonad-sex chromosome scores for each gene in Fig 1H.

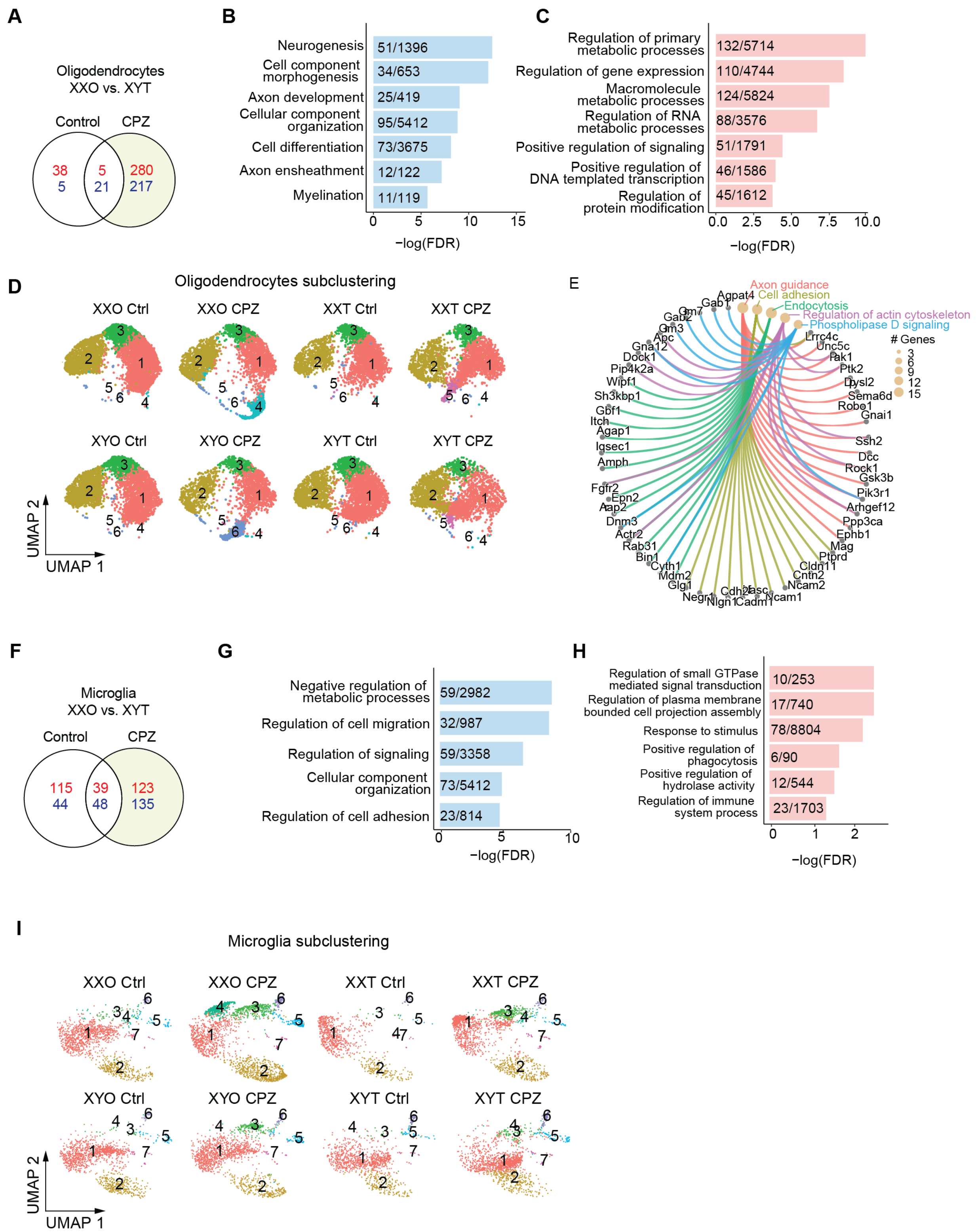


**Figure S4: Sex chromosomes and gonads modify transcriptomes of oligodendrocytes and microglia pre-demyelination.** (A) Venn diagram showing number of significant DEGs in oligodendrocytes between XXO female and XYT male samples in control (n=3) or cuprizone (n=3) states. Red color indicates upregulated genes whereas blue indicates downregulated genes, log2FC cutoff <-0.25 or >0.25. (B) Gene ontology pathway analysis of downregulated oligodendrocyte DEGs between XXO and XYT samples treated with cuprizone (highlighted in green in panel B). Fill numbers indicate number of overlapping genes in dataset/number of total genes in pathway. (C) Gene ontology pathway analysis of upregulated oligodendrocyte DEGs between XXO and XYT samples treated with cuprizone. Fill numbers indicate number of overlapping genes in dataset/number of total genes in pathway. Red color indicates upregulated genes whereas blue indicates downregulated genes. (D) UMAP of 30,011 oligodendrocytes across 6 subclusters split by genotype and treatment. (E) KEGG pathway analysis of Cluster 4 markers with number of overlapping genes represented by circle size. (F) Venn diagram showing number of significant DEGs in microglia between XXO female and XYT male samples in control or cuprizone states, log2FC cutoff <-0.1 or >0.1. (G) Gene ontology pathway analysis of downregulated and (H) upregulated microglia DEGs between XXO and XYT samples treated with cuprizone (highlighted in green in panel B). Fill numbers indicate number of overlapping genes in dataset/number of total genes in pathway. (I) UMAP of 12,473 microglia across 7 subclusters split by genotype and treatment.

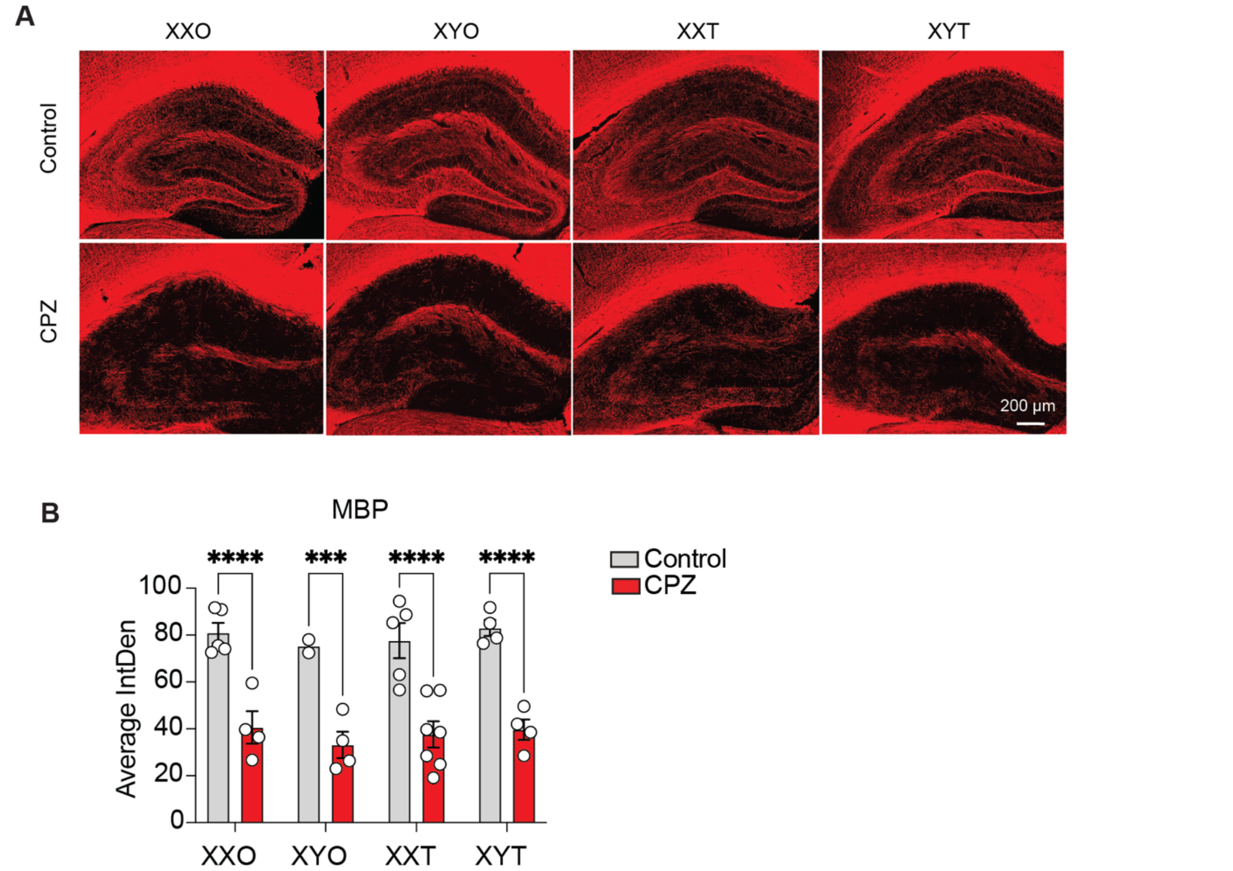


**Figure S5: 5-week cuprizone treatment instills hippocampal demyelination.** (A) Representative images of MBP immunofluorescence in control and cuprizone-treated FCG mice (B) Quantification of MBP intensity. All data are represented as mean +/- s.e.m., for data in (B) significance was determined by mixed model with Tukey's post hoc multiple comparisons test. *$p<0.05$, **$p<0.01$, ***$p<0.001$, ****$p<0.0001$.

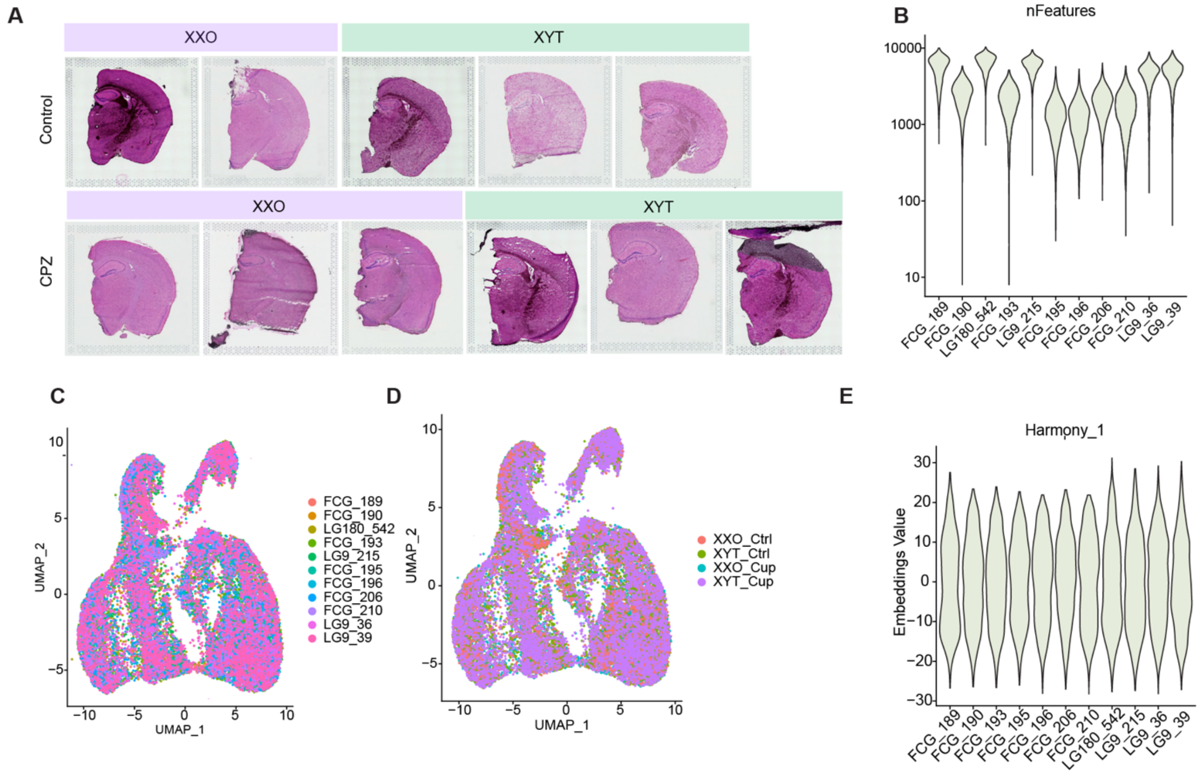


**Figure S6: Quality control assessment of spatial transcriptomics of 5-week CPZ for FCG cohort.** (A) H&E staining of sections in spatial transcriptomics experiment (B) Violin plot showing number of RNA features by sample (C) UMAP showing Visium spots colored by sample (D) UMAP showing Visium spots colored by condition (E) Violin plot showing harmony package component 1 across samples.

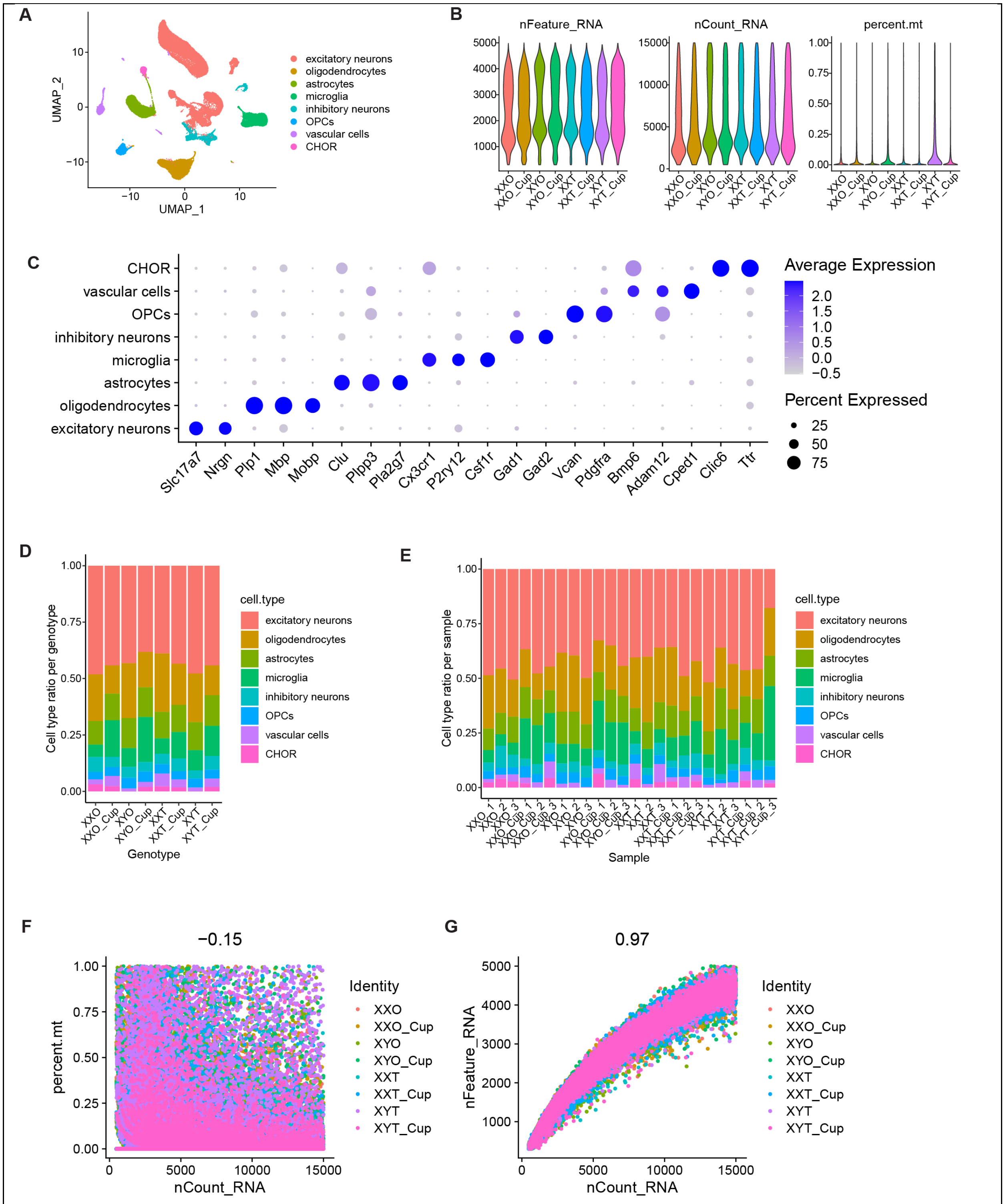

**Figure S7: Quality control assessment of single-nuclei RNA sequencing of 5-week cuprizone for FCG cohort.** (A) UMAP of all nuclei by cell type, n=3 per genotype per treatment. (B) Quality-control plots showing equivalent amounts of total RNA features, total number of RNA counts, and percent mitochondrial RNA in nuclei used for downstream analyses. (C) Summary of genes used for cluster classification into different cell types. (D) Proportion of each cell type detected across the different genotypes. (E) Proportion of each cell type detected across the different samples. (F) Correlation between RNA count and percentage of mitochondrial genes per nuclei and (G) RNA count and RNA features for all samples.

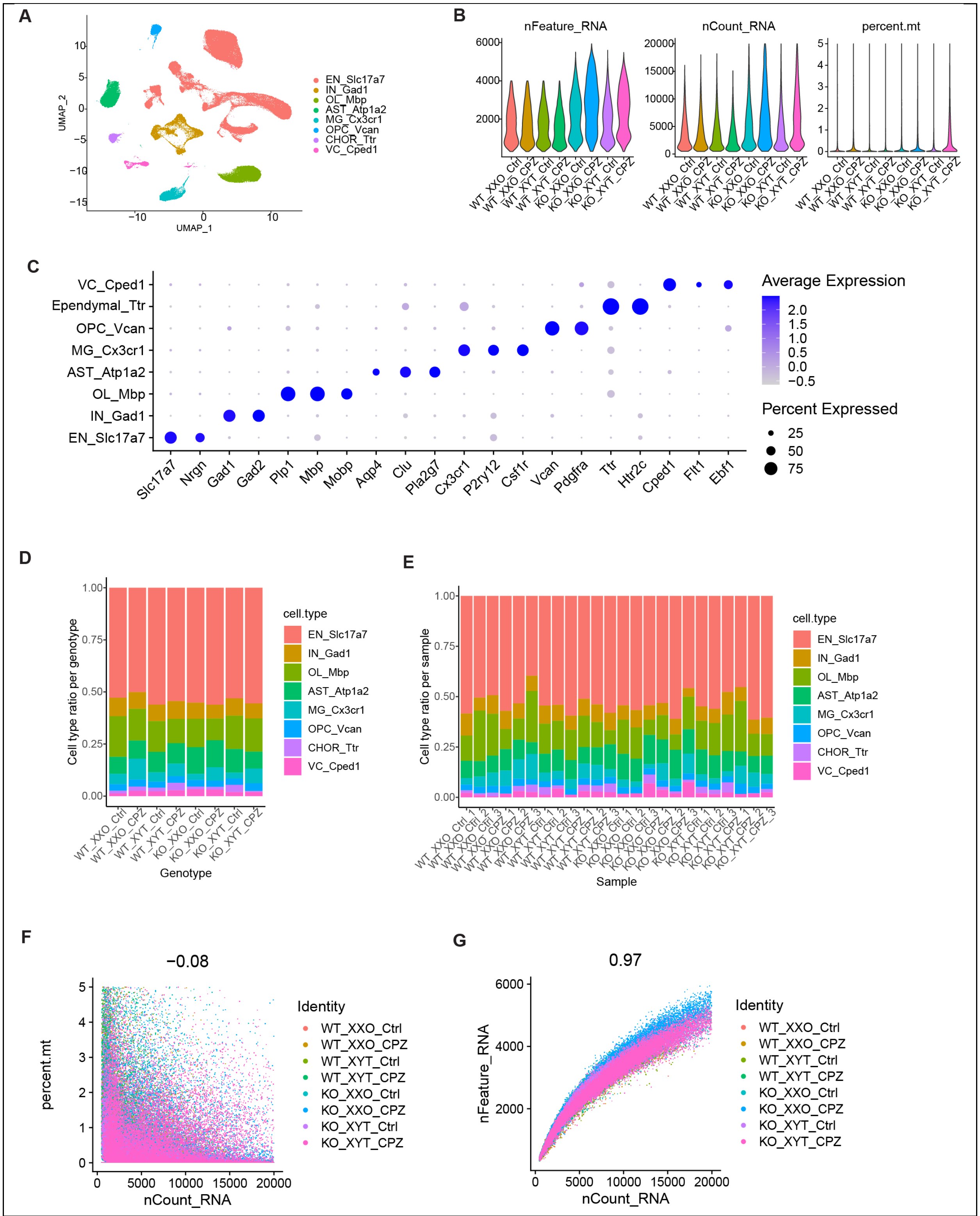


**Figure S8: Quality control assessment of single-nuclei RNA sequencing of 3-week cuprizone for TLR7KO cohort (Related to Fig 6).** (A) UMAP of all nuclei by cell type, n=3 per genotype per treatment (B) Quality-control plots showing equivalent amounts of total RNA features, total number of RNA counts, and percent mitochondrial RNA in nuclei used for downstream analyses. (C) Summary of genes used for cluster classification into different cell types. (D) Proportion of each cell type detected across the different conditions. (E) Proportion of each cell type detected across the different samples. (F) Correlation between RNA count and percentage of mitochondrial genes per nuclei and (G) RNA count and RNA features for all samples.

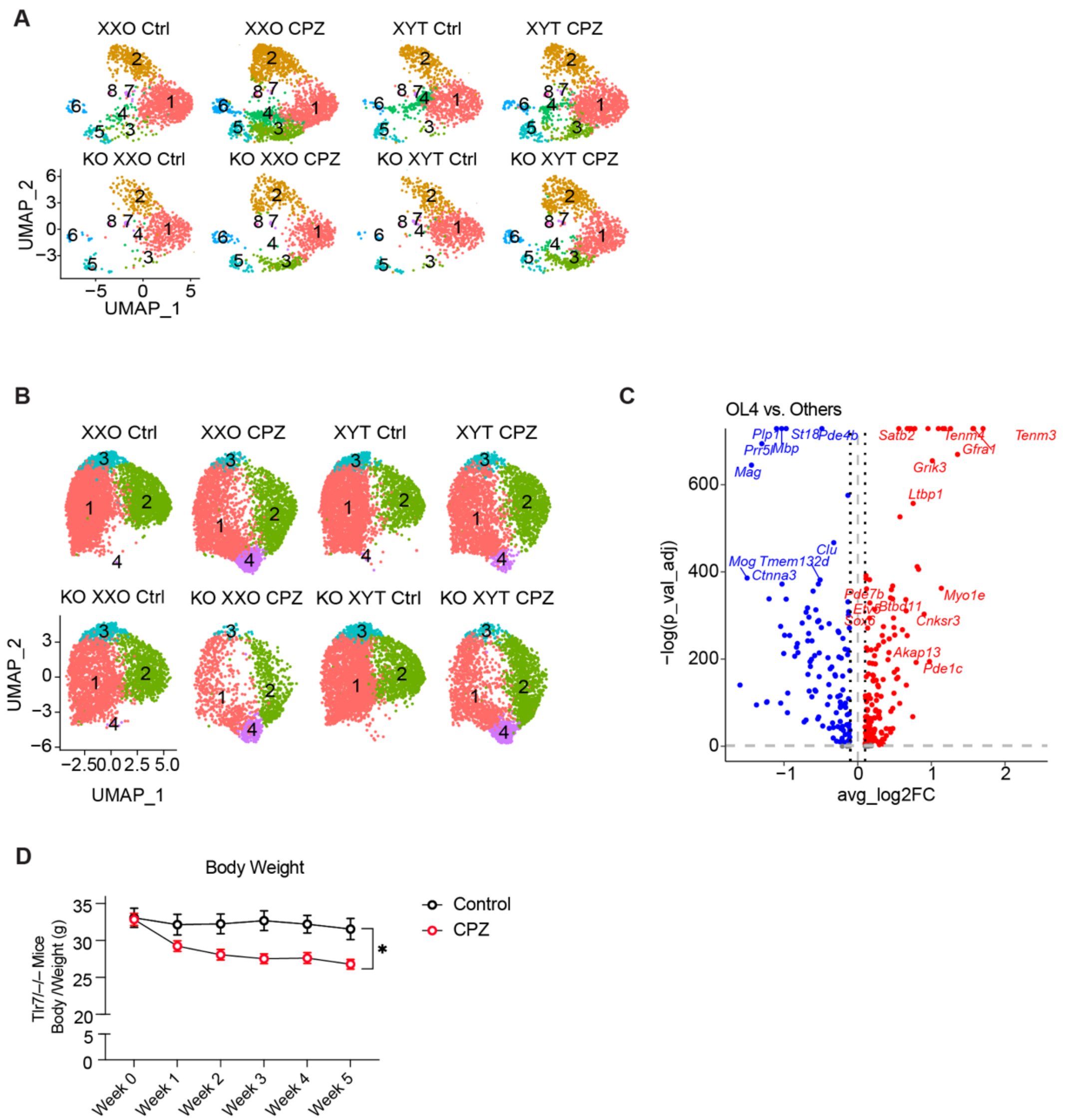


**Figure S9: Microglia and oligodendrocyte subclusters in integration of TLR7KO and FCG single-nuclei RNA sequencing.** (A) UMAP of microglia across 8 subclusters split by genotype and treatment. (B) UMAP of 28,175 oligodendrocytes across 4 subclusters split by genotype and treatment. (C) Volcano plot of DEGs in cluster 4 compared to all other clusters. Dashed lines represent log2foldchange threshold of 0.1 and adjusted p value threshold of 0.05. (D) Weight change in control compared to 5-week cuprizone-treated groups of TLR7-/- mice. All data are represented as mean +/- range, for data in (D) significance was determined by two-way ANOVA with Tukey's post hoc, *p<0.05.

**Supplementary Tables (tables in Excel format)**

**Table S1. Sex-biased gene expression before myelin loss**
Table summarizing snRNA-seq differentially expressed genes across the detected cell types, including X-linked genes by cell type, X-linked genes with sex-biased expression, and the sex chromosome/gonad influence scores.

**Table S2. Oligodendrocyte gene expression pre-demyelination**
Table providing snRNA-seq data for oligodendrocytes, including subcluster markers and differential gene expression of unique cluster 4.

**Table S3. Microglia gene expression pre-demyelination**
Table providing snRNA-seq data for microglia, including subcluster markers and differential gene expression of disease-associated microglia cluster 3 and unique cluster 4.

**Table S4. Neuroinflammatory response at different stages of demyelination**
Table providing multiplex ELISA data at 3 week and 5 week treatment times across hippocampus and frontal cortex lysate.

**Table S5. Spatial transcriptomics post demyelination**
Table providing spatial transcriptomics data, including subcluster markers and genes with sex-biased gene expression across the hippocampal, white matter, and dorsal cortex clusters.

**Table S6. Oligodendrocyte gene signatures at different stages of demyelination**
Table providing snRNA-seq data comparison for oligodendrocytes at 3 week treatment versus 5 week treatment, along with comparison of oligodendrocytes from female and male mice at 5 week treatment, followed by comparison of ovaries versus testes genotypes, and XX versus XY genotypes at 5 weeks of treatment.

**Table S7. Microglia gene signatures at different stages of demyelination**
Table providing snRNA-seq data comparison for microglia at 3 week treatment versus 5 week treatment, along with markers for microglia from 5 week treatment time, differentially expressed genes in cluster 2, and differentially expressed genes in XX versus XY genotypes within cluster 2.

**Table S8. Gene profiles of cell types in vitro myelin treatment and with *Tlr7* knockout**
Table providing bulk RNA-seq data of differentially expressed genes between microglia from male and female mice *in vitro* treated with myelin, comparison of sex-biased genes between in vitro myelin and in vivo cuprizone, and snRNA-seq data from Tlr7 knockout mice treated with cuprizone, including microglia and oligodendrocyte markers.
